# Mechanism of treatment-free remission in patients with chronic myeloid leukemia revealed by a computational model of CML evolution

**DOI:** 10.1101/2022.05.20.492875

**Authors:** Xiulan Lai, Xiaopei Jiao, Haojian Zhang, Jinzhi Lei

**Author notes:** Corresponding author *Email address:* (Jinzhi Lei). These authors contributed equally to this work (Xiulan Lai, Xiaopei Jiao).

## Abstract

In the past few years, international treatment guidelines for chronic myeloid leukemia (CML) have incorporated recommendations for attempting discontinuation of treatment with tyrosine kinase inhibitors (TKIs) outside of the setting of a clinical trial with the aim of treatment-free remission (TFR). Multiple clinical studies have shown consistent results that 40%-50% patients can achieve long-term TFR after TKI discontinuation, and most relapse patients undergo molecular recurrence within 6 months after TKI discontinuation, however the underling mechanisms remain unclear. To understand the mechanism of TFR in patients with CML, we consider the competition between leukemia stem cell and bone marrow microenvironment, and develop a mathematical model to investigate the CML progression dynamics. Model simulations are consistent with clinical observation of CML progression, and reveal a mechanism of dimorphic response after TKI discontinuation. Our model predicts that neoplasitic microenvironment is significant for CML occurrence and progression. We furthermore discuss the prediction of TFR based on the change rates of microenvironment index and leukemia stem cells ratio prior TKI discontinuation.

## Introduction

Chronic myeloid leukemia (CML) is a model disease for targeted therapy, characterized by the Philadelphia chromosome and its molecular counterpart, the BCR-ABL1 fusion gene, encoding a tyrosine kinase with aberrant activity [1, 2]. In the past decade, tyrosine kinase inhibitors (TKIs) have been the standard treatment and improve the survival of CML patients [3, 4, 5]. More than 85% of CML patients treated with TKIs achieve a complete cytogenetic response and approximately 40% of these patients achieve a complete molecular remission (CMR) [6, 7]. Many patients with deep molecular responses have a prerequisite for TKI therapy discontinuation with the aim of treatment-free remission (TFR) [8, 9]. Many clinical trial experience of TFR have been studied over the years, however, the concept of TFR is relatively new, and little is known about the molecular components regulating TFR [10].

Multiple clinical studies have investigated TFR after the first-generation TKI imatinib therapy [11, 12, 13, 14, 15]. All studies have near-identical inclusion criteria and come out with consistent results. After stopping imatinib in patients with stable undetectable BCR-ABL1 for at least 2 years, the overall molecular relapse rate was about 60%, and long-term molecular survival was about 40%. Molecular recurrence (MR) occurred mostly within 6 month after imatinib stopping, and late relapse were rarely observed. In a long-term follow up of the French stop imatinib (STIM1) study [14], molecular recurrence-free survival (RFS) was 43% (95% CI, 33% to 52%) at 6 months and 38% (94% CI, 29% to 47%) at 60 months. Similar results are also seen in studies of second-generation TKIs [16, 17, 18, 19], with a 96-weak molecular RFS of 49% in the ENESTfreedom study [18] and 53% in the ENESTop study [19], both were designed to evaluate the effects of nilotinib discontinuation. Moreover, in either first- or second-generation studies, most relapsed patients who reinitiated original TKI regained major molecular response (MMR) (BCR-ABL1 ≤ 0.1%). Some patients can reach second TFR after the first failed attempt and restart TKI therapy [20].

In clinical studies, CML patients starting TKI discontinuation from near-identical conditions show different outcomes, either relapsed within 6 months, or achieve long-term TFR (with rare late relapsed cases). It is well known that TKIs treatment is unable to completely eradicate leukemic stem cells (LSCs), residual BCR-ABL1^+^ cells can be observed during TFR [18, 21]. There must be some other conditions to maintain the long-term TFR. Multiple parameters may associate with TFR in individual cases, such as age and sex [11, 22], Sokal score [11], duration of TKI treatment [11, 12, 23, 24], or the immune status [16, 17, 25, 26]. The interplay between leukemia cells and the bone marrow microenvironment (BMM) is important for CML progression [10, 27, 28]. Optimal predictors of successful TFR are yet to be defined [8].

The dynamics of leukemic-related hematopoietic system is essential for understanding the clinical outcome of CML. Mathematical modeling of relapsing CML through model assumption from stem cell biology predicts a rapid increase in BCR-ABL1 when treatment is withdrawn [29, 30, 31], and TFR would not be predicted without the stem cell-niche interaction. Hence, additional interactions have to be involved in order to explain the outcome of TFR. Here, we investigate a mathematical model of hematopoiesis that describes both healthy and leukemia stem cells (LSCs), as well as the leukemia-microenvironment crosstalk. The level of leukemia in patients is quantified by measuring the leukemia/healthy cells ratio in circulating blood. Imatinib therapy is incorporated into the model as to increase the death rate of leukemia cells. By comparing model dynamics with clinical data, we identify model parameters to reproduce the three stages of CML development. Furthermore, model simulations allow us to investigate the dynamics of patient responses after imatinib discontinuation, and provide evidence of how the leukemia-microenvironment crosstalk play essential role in predicting TFR of individual patients.

## Results

### Mathematical model of CML dynamics

To model the CML evolution process, we considered the dynamics of hematopoietic stem progenitor cells (HSPCs) and leukemia stem progenitor cells (LSPCs) in BMM, and the interaction between leukemia cells with surrounding niche to render the variance between normal microenvironment (NME) and tumor microenvironment (TME). Population dynamics of hematopoietic cells in peripheral blood (PBHC) and leukemic cells in peripheral blood (PBLC) were also included in order to compare with clinical criteria. The schematic representation of the interactions is shown in Fig. 1. Detailed mathematical model formulations are given in Materials and Methods.

**Figure 1:**
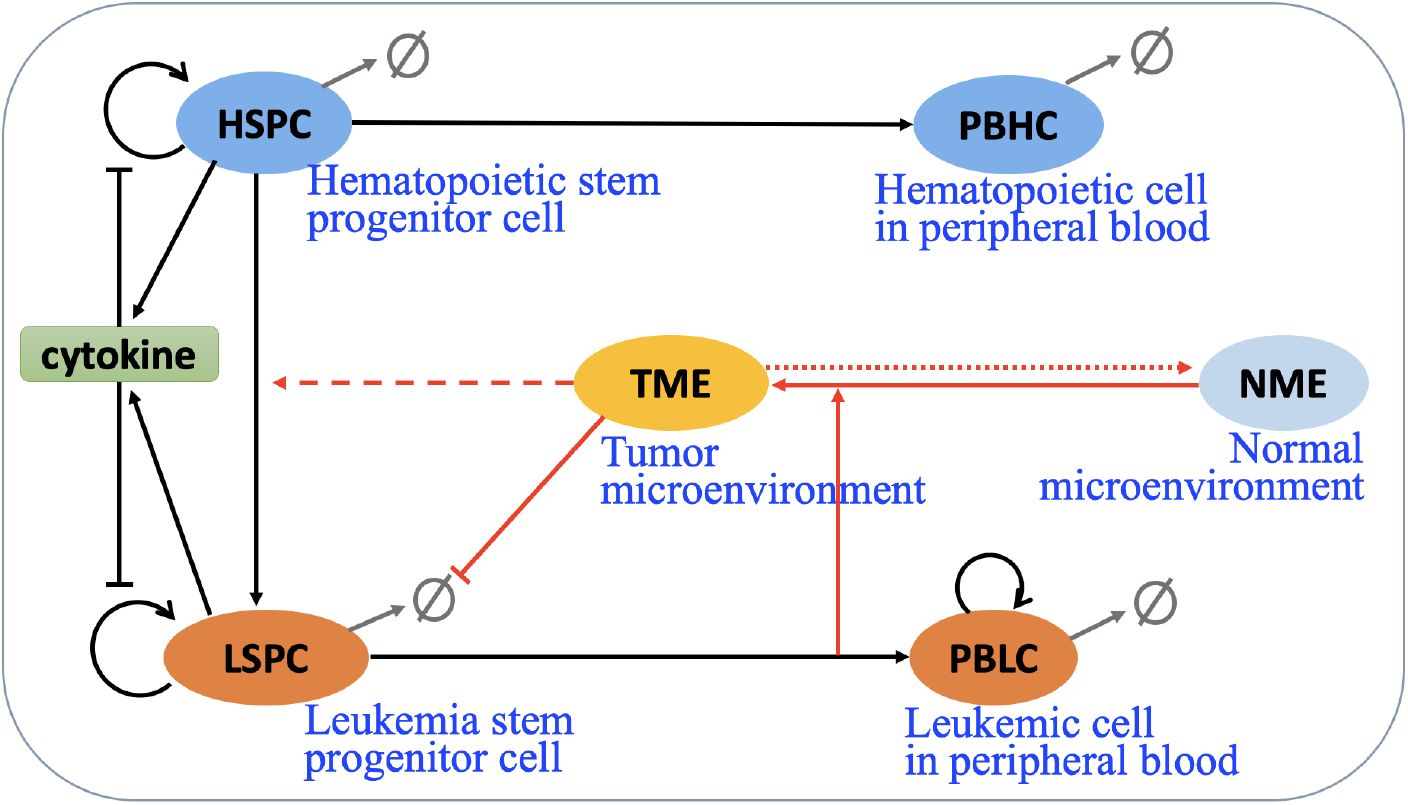
Schematic representation of CML evolution driven by tumor microenvironment. Black arrows indicate cell interaction/transitions, red arrows show the interactions with microenvironment, grey arrows represent cell death.

Leukemia cells are known to interact with their surrounding niche in order to render it more permissible for leukemia progression [32, 33, 34, 27]. Detailed molecular mechanisms of leukemia-BMM interactions were omitted in the model. However, we introduced a variable *Q* to represent the microenvironmental condition that can change between NME and TME, and leukemia cells can induce the transition from NME to TME. We further assumed that TME promotes the survival of leukemia cells and the potential frequency of generating leukemia cells from normal HSPC to LSPC. Interactions for the microenvironment are shown by red arrows in Fig. 1.

### Model validation with CML evolution

Clinically, CML progression includes three phases characterized by the BCR-ABL1 level in peripheral blood, namely chronic phase (CP), accelerated phase (AP) and blast crisis (BC), with BCR-ABL1 levels < 10%, 10% ∼ 20%, and beyond 20%, respectively [35]. To validate the proposed model, we identified model parameters and compared the simulated CML progression with clinical data.

To quantify the CML evolution, we analyzed gene expression changes in 91 cases of CML in chronic (42 cases), accelerated (17 cases) and blast phases (32 cases) [36]. CD34 expression showed positive association with CML progression, and a CD34^+^ similarity score was proposed to represent the disease progression marker [37]. To compare the disease progression of different patients and compare with model simulation, we defined the CML progression age starting from the emerge of BCR-ABL1 cells through CD34 gene expression level (see Fig S1 and (4) in Materials and Methods), and tracked disease progression stages of mixed-population clinical samples through blast count versus the CML age (Fig. 2A).

**Figure 2:**
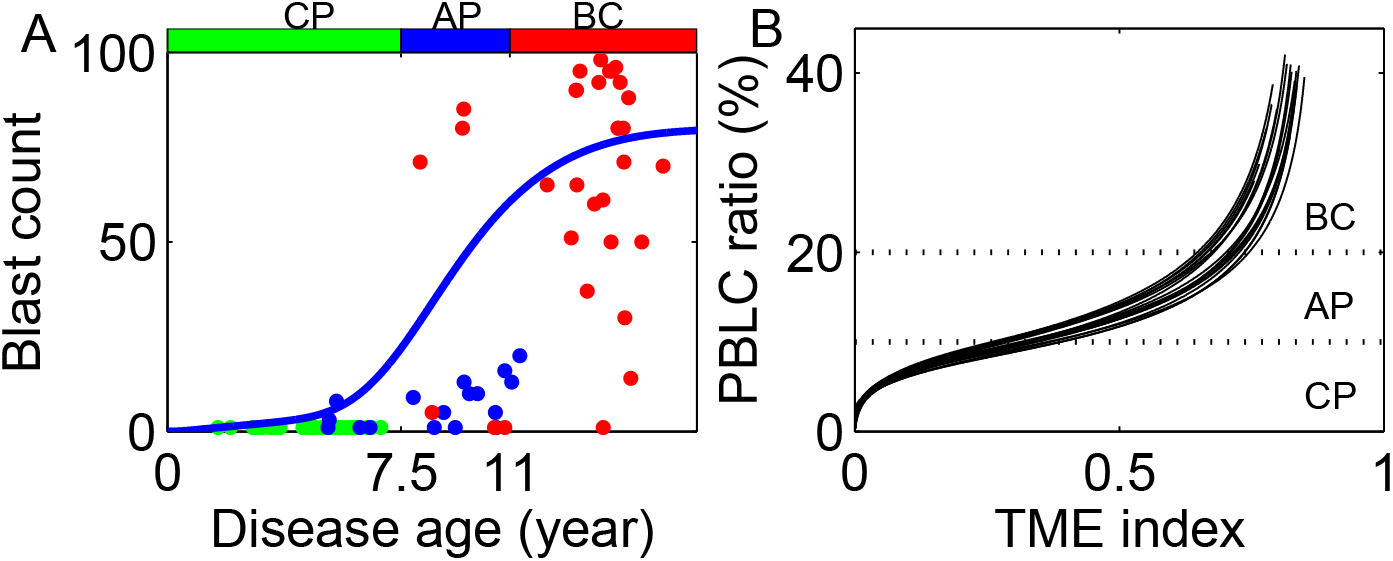
CML evolution. (A) Blast count evolution. Green, blue and red dots denote CP, AP and BC patients, respectively. Red curve represents numerical results of our model. (B) PBLC percentage in peripheral blood versus TME index during CML evolution without therapy.

Blast count represents the concentration of leukemia cells in peripheral blood, which is proportional to numbers of PBLC in the model. Fig 2A shows the simulated evolution of PBLC after the occurrence of the first LSPC. The average dynamics of multiple sample simulations show three stages dynamics in the blast counts, which include slow development during CP, fast increase during AP, and high level maintenance during BC. Specifically, the three stages progression is characterized with the combination of the BCR-ABL1 ratio *r*_PBLC_ in peripheral blood and the TME index (Fig. 2B). During the CP phase, TME index increases rapidly, with slow increase in the ratio *r*_PBLC_. In the AP phase, *r*_PBLC_ accelerated increase, with slowdown of TME index changes. In the BC phases, the TME index is mostly unchanged and *r*_PBLC_ rapidly increase to reach a saturation level.

### CML progression with imatinib therapy

The survival of CML patients were very low before 1975 (no treatment), and has significantly and obviously improved since 2001 due to the administration of TKI therapy [38]. A randomized CML-study after a long-term evaluation of imatinib shown that 10-years overall survival has reached 85% - 90% [39]. To compare simulation results with clinical survival data, we assumed a probability of mortality that is dependent on the BCR-ABL1 ratio *r*_PBLC_, and tuned the model parameters to fit the survival curve for patients without treatment (before 1975) based on clinical data from MD Anderson Cancer Center [38] (Fig. 3A).

**Figure 3:**
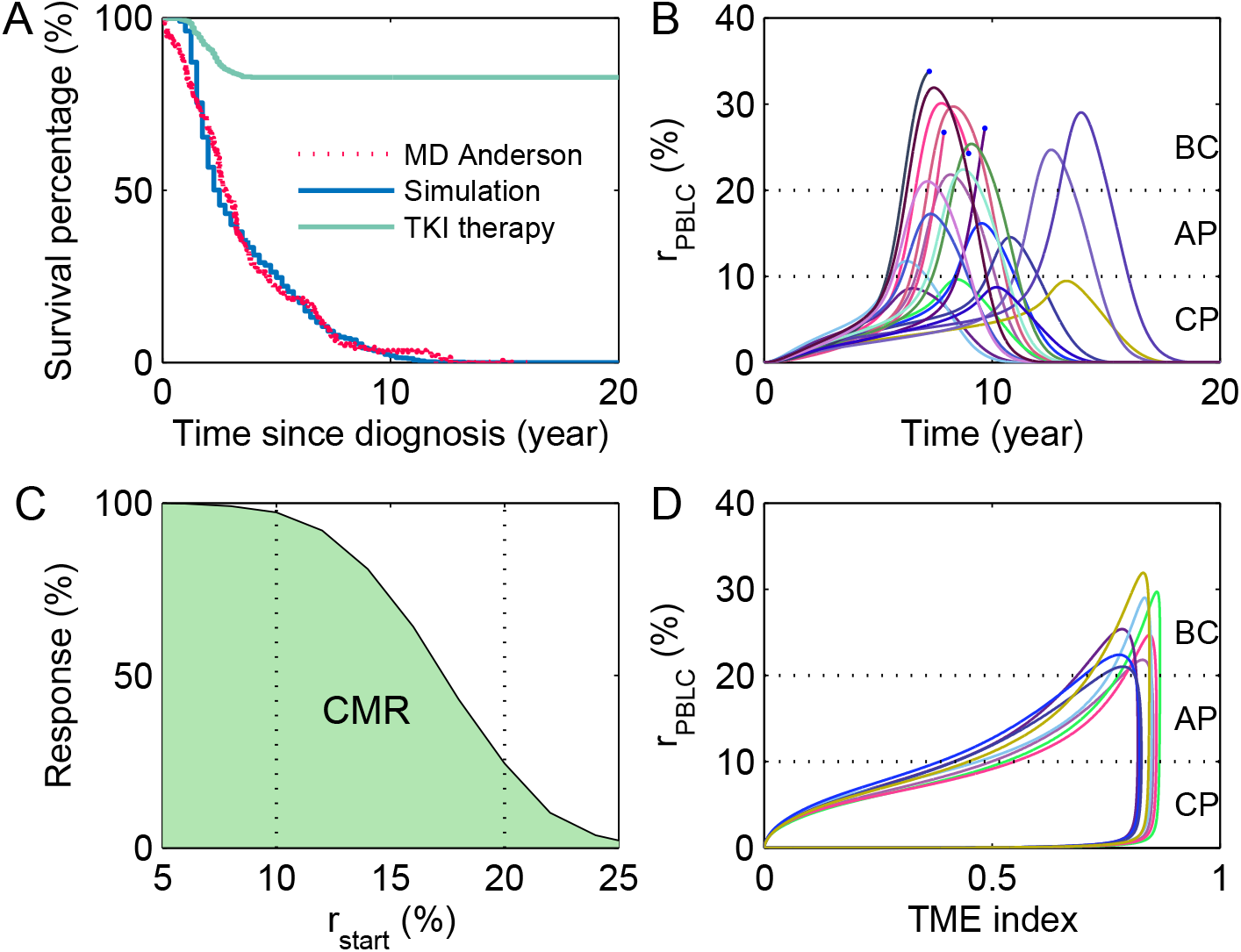
CML evolution for patients under continuous imatinib treatment. (A) Overall survival of CML patients. Red dashed curve for MD Anderson Cancer Center data for patients without treatment [38]; Blue curve for model simulation of patients without treatment; Green curve for model simulation of patients with continuous imatinib treatment. (B) The dynamics of PBLC ratio (*r*_PBLC_) for 20 patients with continuous imatinib therapy. Here, the imatinib therapy was assumed to start for a patient randomly after the PBLC ratio reaches a level between 5% − 20%. (C) The percentage of CMR in CML under different starting ratio of PBLC in peripheral blood (*r*_start_). (D) The relation of PBLC ratio and TME index along CML evolution prior therapy and under continuous imatinib treatment. All parameter values are the same as in Tables 1 and 2 in Materials and Methods.

Next, to model the effects of imatinib therapy, we introduced an extra leukemia cell death rate that is dependent on the drug dose (see (2) in Material and Methods), and adjusted model parameters to reproduce the overall long-term survival of 80% - 90% after continuous TKI treatment (Fig. 3A). In simulations, virtual patients were first generated to optimize the overall survival without treatment, and next each patient was assumed to be diagnosed and start TKI therapy randomly when the ratio of leukemia cells in peripheral blood (*r*_PBLC_) is in the range 5% − 20%. Fig. 3B shows the time evolution of *r*_PBLC_ of 20 sample patients under continuous TKI treatment. Results showed that *r*_PBLC_ may increase shortly after the onset of TKI treatment, followed by continuous decrease in most patients and develop to CMR as the leukemia cell ratio reduces to an undetectable level; however, a few patients of late treatment can still develop to mortality. Simulations showed that the initiation timing of starting TKI treatment significantly influences the potential of CMR, patients diagnosed and treated earlier (with low PBLC proportion) have much higher probability of long-term survival, and there is a significant decline in the survival rate for patients diagnosed in the accelerated phase (Fig. 3C).

The CML progression to CMR after imatinib therapy is further explored through the combination of BCR-ABL1 ratio *r*_PBLC_ and the TME index (Fig. 3D). During the recovery process, *r*_PBLC_ rapidly decreases from BC towards AP with mostly unchanged in the TME index. In the CP phase when *r*_PBLC_ is very low (*r*_PBLC_ < 1%), the TME index decrease toward a NME, and slow decreasing of *r*_PBLC_ toward the final complete remission. We noted the difference between CML evolution and the remission process, and the TME index can be different for the same level of BCR-ABL1 ratio in the two processes during the CP phase. We asked whether microenvironment condition is essential for TFR after imatinib discontinuation.

### CML progression after imatinib discontinuation

Now, we investigate CML progression after imatinib discontinuation by setting the drug dose as zero and continue the model simulation for at least 15 years. Here, we stop treatment when the PBLC ratio reaches 0.01% (*r*_stop_ = 0.01%). CML progression of virtual patients are shown in Fig. 4A, in which some patients show leukemia cell recurrence shortly after TKI discontinuation, while other patients show long-term TFR with extreme low PBLC ratio in peripheral blood. For the TFR patient, the PBLC ratio slightly increases after treatment discontinuation, and reversibly decreases toward an undetectable level (Fig. 4A).

**Figure 4:**
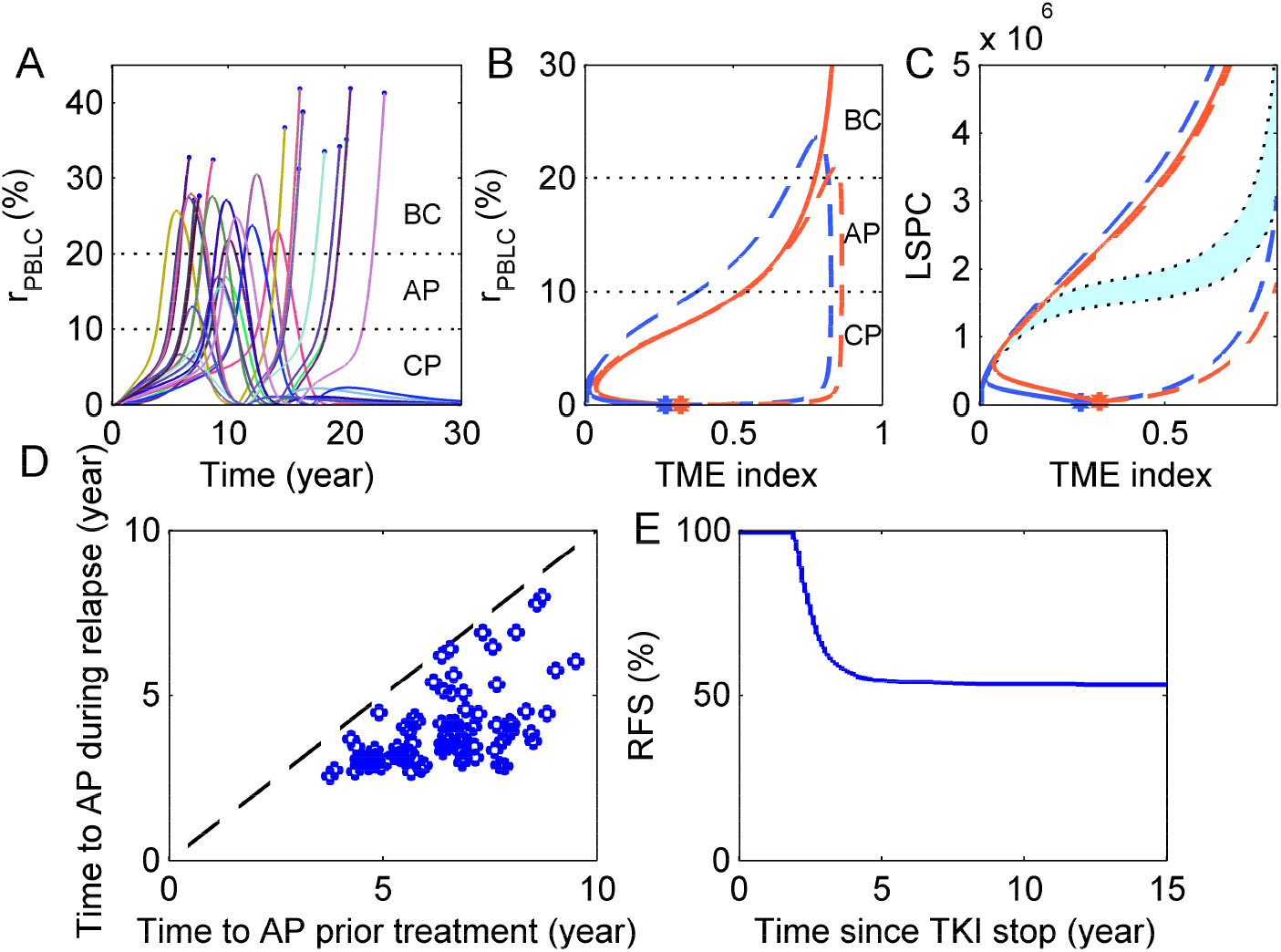
CML evolution for patients with discontinuous TKI therapy. (A) Dynamics of PBLC ratio (*r*_PBLC_) for 20 patients with TKI therapy initiated randomly when 5% < *r*_PBLC_ < 20% and stopped when *r*_PBLC_ = 0.01%. (B) Typical trajectories of PBLC ratio versus TME index for two patients with relapse (red) and TFR (blue) after treatment discontinuation. Dashed lines and solid lines indicate the trajectories before and after treatment discontinuation, respectively, while the star points show the points of treatment discontinuation. (C) Trajectories of LSPC versus TME index for the two patients in (C) with relapse (red) and TFR (blue), which is enlarged around the time point of treatment discontinuation (red and blue star points). Cyan zone shows the critical values of LSPC and TME index that separate the fate of either tumor relapse or TFR(see Materials and Methods). (D) Comparison of the time to AP during phrases of prior treatment and during relapse process after stopping treatment in each relapsed patient. (E) Molecular relapse-free survival (RFS) curve after TKI stop. All parameter values are the same as in Tables S1 and S2.

Typical dynamics for relapsed and TFR patients are shown in Fig. 4B, respectively. At the time of treatment discontinuation (red and blue star points), the PBLC ratios are very low for both cases. However, the relapsed patient (in red) showed rapid increase in *r*_PBLC_ and TME index, and further developed to the irreversible AP phase, while the TFR patient (in blue) showed initial increase then rapid decline of *r*_PBLC_ and continuous decrease of TME index. There is a zone of critical values for LSPC number and TME index as shown in Fig. 4C (cyan zone), when a patient trajectory passes across this zone after treatment discontinuation, the LSPC number and TME index would increase rapidly and leading to tumor relapse, otherwise, LSPC number and TME index would decline whereby reaches TFR. The critical zone is determined by the dynamics of how TME index is dependent on LSPC levels. We compared the time periods developed to AP phase in early disease progression and in the relapse stages for each relapsed patients, the periods in the relapse process are much shorter than that in the early progression stage (Fig. 4D). This result indicates that relapse process developed much faster than the early disease progression.

The above typical dynamics suggests a threshold of PBLC ratio as a criterion of leukemia cells recurrence. Based on the defined threshold, the molecular relapse-free survival (RFS) curve was obtained from a cohort of 1000 virtual patients who stop treatment at MR4.0 (PBLC ratio *r*_PBLC_ < 0.01%) (see Fig. 4E). Here, molecular relapse was defined as *r*_PNLC_ > 5%, and hence there is a lag time of about 2 years of PFS decreasing after treatment stop. The simulation result is in agreement with clinical trials, with about 47% patients rapidly develop to molecular recurrence after treatment discontinuation. There is a clear plateau in the molecular RFS curve, which indicates the outcome of long-term TFR. Thus, both clinical study and modeling simulations support the dichotomous patients of either early molecular relapse or TFR after TKI treatment.

### Mechanisms of TFR after imatinib discontinuation

To explore the factors associated with molecular relapse after imatinib discontinuation, we investigated virtual patient responses under various conditions prior to imatinib discontinuation. First, we varied the PBLC ratio at the time of treatment stop (*r*_stop_), which is crucial to the molecular RFS survival. Higher level of the ratio *r*_stop_ obviously increase the potential of early molecular relapse and reduce the probability of TFR (see Fig. 5A). The total duration of imatinib treatment and the duration of DMR (MD4.0 or lower) prior to imatinib discontinuation are known to associated with a higher probability of TFR [8]. Model simulations showed that the duration of imatinib treatment is significant for the outcome of treatment discontinuation, early stop (≤ 5 years) may result in high relapse probability (> 90%) and should be against, the cumulative relapse probability can reduce to 50% after 6 years treatment, and 20% after 10 years treatment (see Fig. 5B). Extension of the MDR duration is positively related to the probability of TFR; the probability can raise to above 70% with 2 years more treatment during DMR prior to imatinib discontinuation (see Fig. 5C).

**Figure 5:**
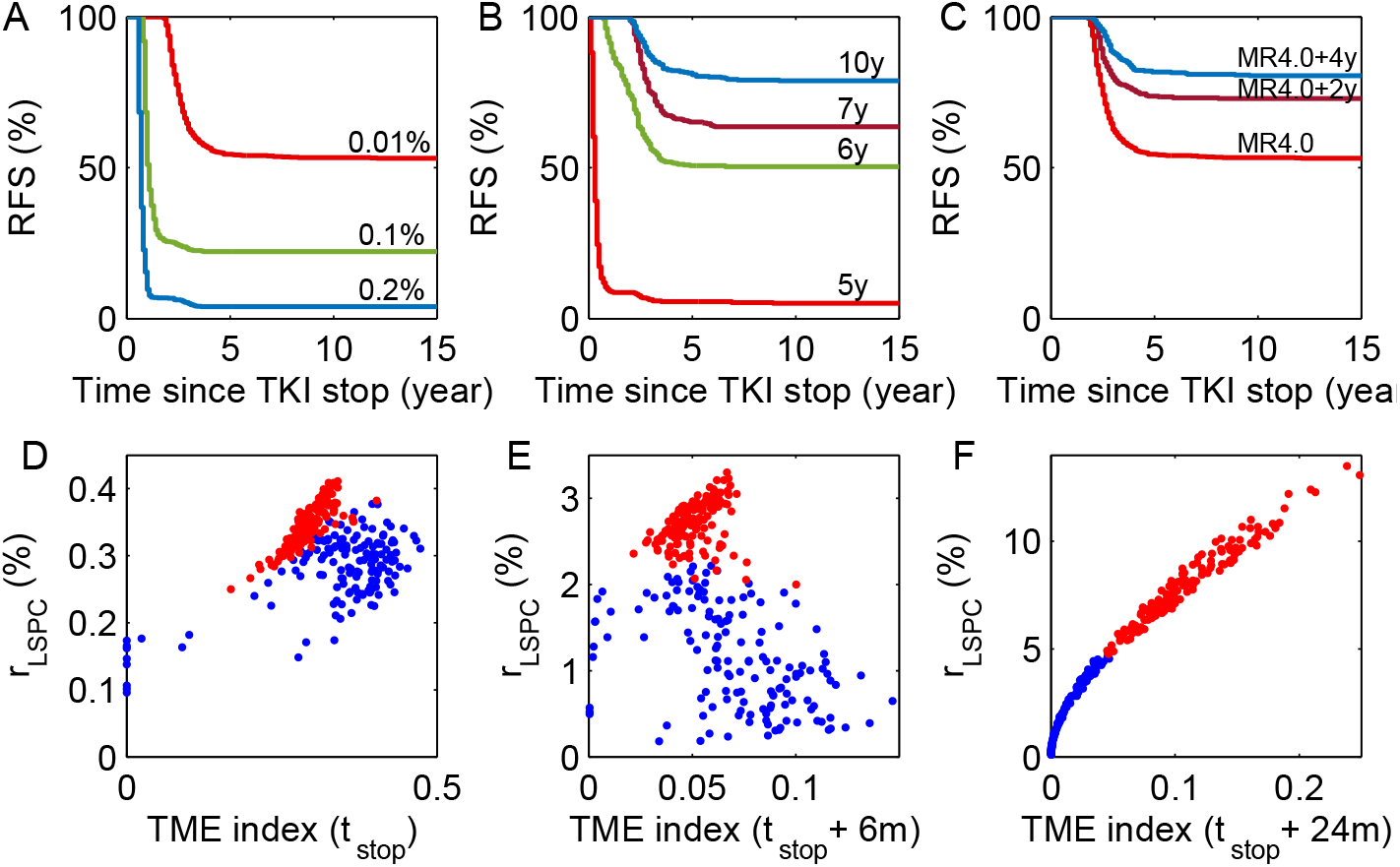
Mechanisms of TFR after imatinib discontinuation. (A) Molecular RFS curves of patients stopped imatinib therapy at PBLC percentage level, *r*_PBLC_ = 0.01%, 0.1%, 0.2%, respectively. (B) Molecular RFS curves for patients stopped imatinib therapy after treatment for 5, 6, 7, and 10 years, respectively. (C) Molecular RFS curves of patients stopped imatinib therapy when PBLC percentage level reaches *r*_PBLC_ = 0.01% (MR4.0), or with 2 or 4 more years treatment after MR4.0. (D)-(F) LSPC percentage and TME index at the time of stopping treatment, 6 months after stopping treatment and 24 months after stopping treatment, respectively. Here the treatment is stopped when PBLC percentage level reaches *r*_PBLC_ = 0.01%. Red dots for relapsed patients, and blue dots for TFR patients after treatment discontinuations.

To further consider the mechanisms of TFR after imatinib discontinuation, we grouped the virtual patients into either relapsed or long-term TFR, and examined the combination of bone marrow leukemia cells ratio and the TME index at the time of treatment stop. We note that many patients who developed to TFR have nonzero leukemia cells in bone marrow, and the leukemia cells ratio is usually higher in relapsed patients than in TFR patients (see Fig. 5D). In a short period after the treatment stop, the bone marrow leukemia cells ratio continuously increases while the TME index decreases initially for both relapsed patients and TFR patients (see Fig. 5E). However, in a long run, both bone marrow leukemia cells ratio and the TME index increase for the relapsed patients, whereas TRF patients maintain both low leukemia cell ratio in the bone marrow and small TME indices (see Fig. 5F).

In the model, we introduce a function *η* to represent the external factors that may affect the potential frequency of leukemia cells production from normal HSPC. This external effect is necessary to model the initiation of leukemia cells, and could have influence on the leukemia recurrence rate after treatment withdraw. Nevertheless, while we choose a certain pattern of mutation frequency so that *η* becomes zero at the time of treatment withdraw (see Fig. S2A), simulation results show similar dynamics of TME and bone marrow leukemia cells ratio for patients with either molecular relapse or TFR (see Fig. S2B-D). Hence, the outcome of either leukemia recurrence or TFR after treatment withdraw may not rely on the external factors, and interactions between microenvironment condition and leukemia cells can be crucial.

### Prediction of TFR

The combination of bone marrow leukemia cells ratio (*r*_LSPC_) and TME index after treatment stop is essential for TFR. To further identify the predictive markers, we examined the combination of *r*_LSPC_ and TME index in two group of patients at 6m, 12m and 24m prior imatinib discontinuation (see Fig. 6A). Results showed that data collection at 6-12 months prior treatment stop perform well in separating the two groups patients. Prior to treatment stop, both TME and *r*_LSPC_ are decreased, and the relapsed patients showed a more rapid decline of TME and *r*_LSPC_ than those of TFR patients. Comparing the change of TME and *r*_LSPC_ during the period from 2 years prior treatment stop to 2 years after treatment stop, we found that the change rates of TME and *r*_LSPC_ could potentially be served as a predictor of TFR outcome (see Fig. 6B and Fig. 7). The declining rate of TME for relapsed patients is always higher than that of TFR patients prior TKI stop, while after the treatment stop, relapsed patients show a lower declining rate of TME for the first year, and the TME start to increase after the first year, whereas the TFR patients showed higher declining rate of TME for the first year and no raise of TME latter on (see Fig. 7 and Fig. S3). Relapsed patients also show the characteristic of lower *r*_LSPC_ declining rate prior to the treatment stop and high *r*_LSPC_ increasing rate after the treatment stop (see Fig. 7 and Fig. S4).

**Figure 6:**
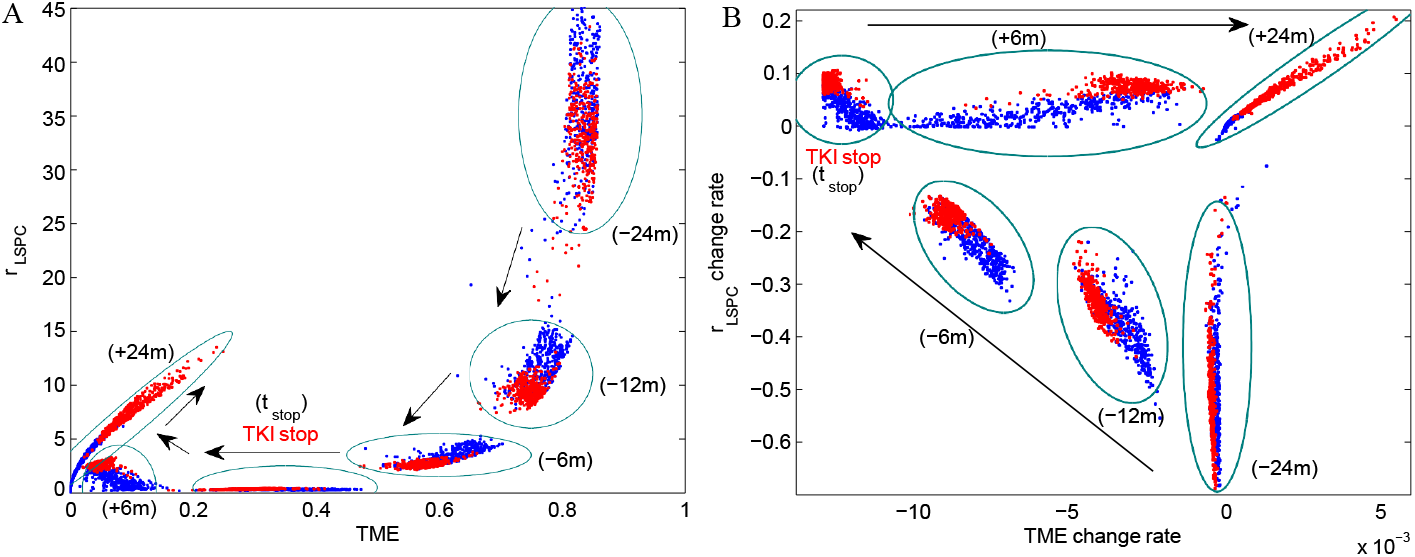
Evolution of TME, bone marrow leukemia cells ratio (*r*_LSPC_) and their change rates for TFR patients and relapsed patients. (A) Evolution of TME and bone marrow leukemia cells ratio for TFR patients (blue dots) and relapsed patients (red dots) at 2 years (−24m), 1 year (− 12m), 6 months (− 6m) before TKI stop, at the time of TKI stop (*t*_stop_), at 6 months (+6m) and 2 years (+24m) after TKI stop. (B) Evolution of change rate of TME and leukemia cell ratio in bone marrow for TFR patients (blue dots) and relapsed patients (red dots) at the time points same as (A). Different time points are shown by green circles. The arrows indicate the direction of time evolution.

**Figure 7:**
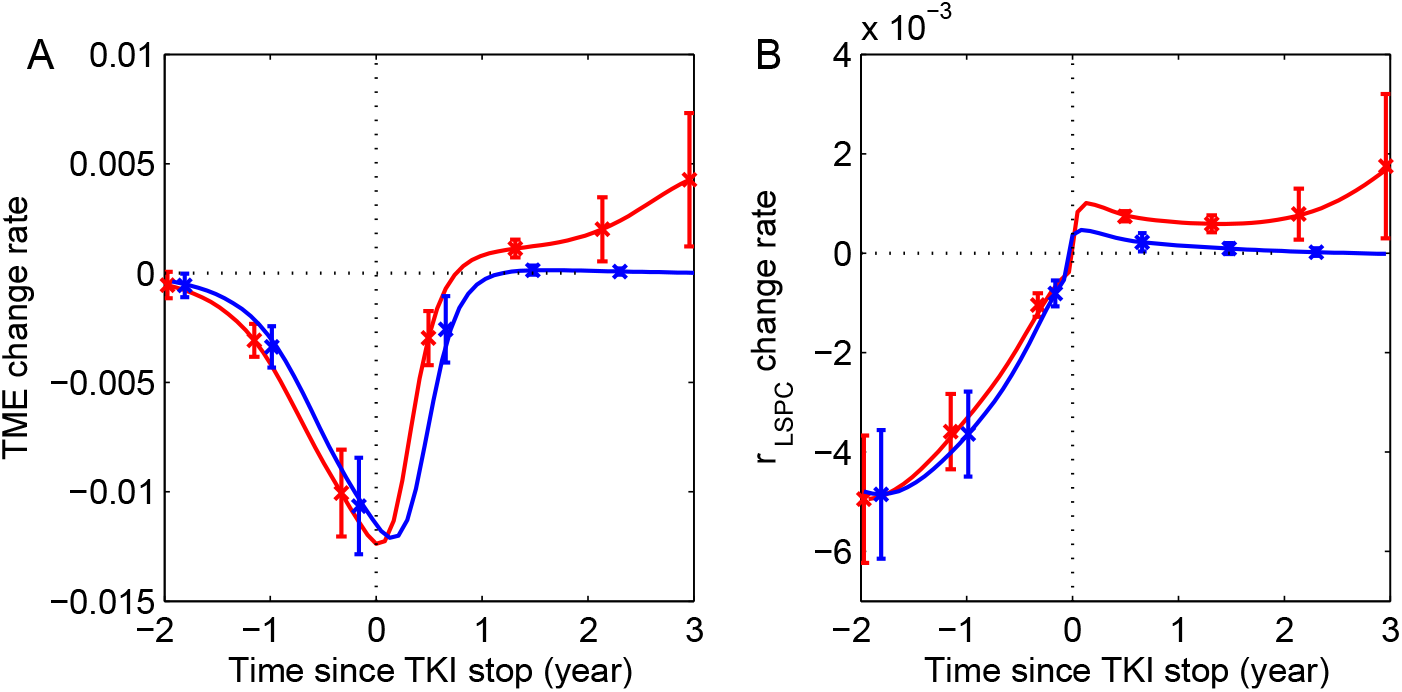
Change rates of TME and bone marrow leukemia cells ratio (*r*_LSPC_). (A) Average change rates of TME index from 2 years prior to TKI stop to 3 years after TKI stop. 0 indicates the time point of TKI stop. (B) Average change rates of bone marrow leukemia cells ratio from 2 years prior to TKI stop to 3 years after TKI stop. Here, the change rates are calculated by the variances in 7 days.

### TFR related parameters

We further examine the parameters that are related to the TFR responses. To this end, we analyzed the parameters for the two group patients of either early relapse or TFR (see Fig. S5). Results show that the two group patients have different preference in the kinetic parameters for the stem cell population, the relapsed patients have higher probability of possessing larger LSPC proliferation rate (*β*_L_) and lower apoptotic rate (*µ*_L_) and differentiation rate (*κ*_L_). Moreover, relapsed patients tends to have smaller transition rate from NME to TME (*κ*_Q_). There is not significant difference in the PBLC ratio to start treatment and the treatment period before treatment stop in the two group patients.

Next, we performed sensitivity analysis for the key dynamic parameters. Results show that continuous TKI treatment time to reach TFR is strongly positively correlated with the LSPC proliferation rate (*β*_L_) and strongly negatively correlates with the proliferation rate of HSPC (*β*_H_ and *θ*_H_) and the LSPC differentiation rate (*κ*_L_) (Fig. S6A). The treatment time to TFR also shows positive correlation with the transition rate from NME to TME (*κ*_Q_) and the late reduction of BCR-ABL1 occurrence rate (*µ*_*η*2_), and negative relation with transition rate from TME to NME (*κ*_I_). Similarly, the length of survival time for the patients who undergo death is strongly positively correlated with the HSPC proliferation rate (*θ*_H_ and *θ*_H_) and negatively correlated with LSPC proliferation rate (*θ*_L_) (Fig. S6B). The transition rate from NME to TME (*κ*_Q_) shows negative correlation with the survival time.

### Second TFR attempt

To investigate the probability of second TFR attempt for patients with molecular recurrence at the first TKI discontinuation, we restarted the treatment when *r*_PBLC_ ≥ 5% and stopped the treatment for second time. The time evolution of *r*_PBLC_ is shown in Fig. 8A. During the second TKI treatment, some patients relapsed after the first TKI discontinuation achieved TFR, which are shown by cyan dots in Fig. 8C at the time of the first TKI stop and in 8D at the time of the second TKI stop. There are some patients with high ratio of LSPC at both time of the first and second treatment stops, who show tumor relapse after the second TKI (red dots in Figs. 8C-D). The RFS rate after treatment stop is elevated significantly after the second treatment in contrast with the first treatment (see Fig. 8B). In our simulation, the relapse rate reduced from 47% at the first TKI discontinuation to about 10% after the second TKI discontinuation. The extension of the second treatment duration further increases the probability of TFR.

**Figure 8:**
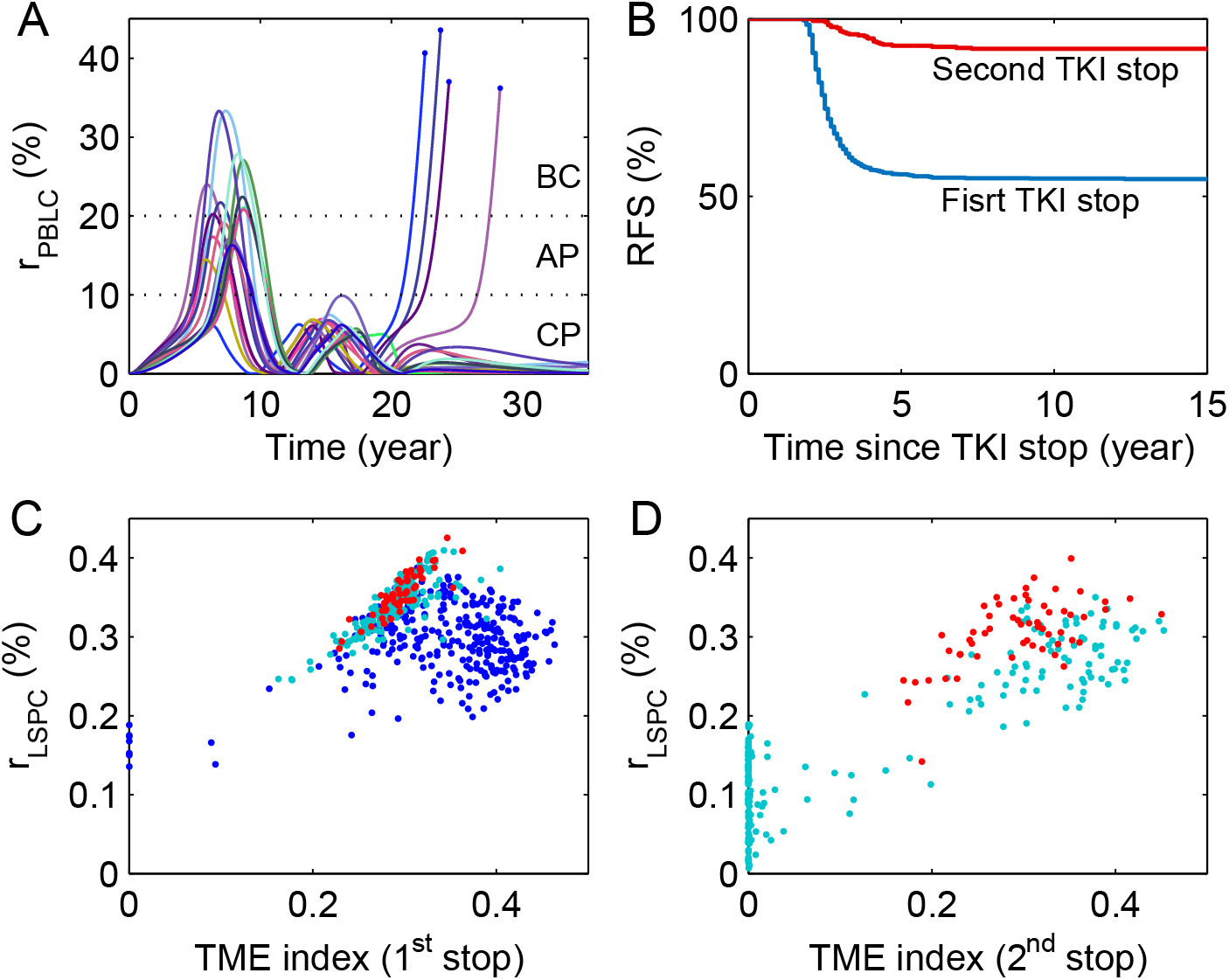
CML evolution and mechanisms of TFR after second imatinib treatment discontinuation. In all the simulations, patients start imatinib treatment at the first time randomly when 5% < *r*_PBLC_ < 20%, stop imatinib treatment at the first time at PBLC percentage level *r*_stop_ = 0.01%, and restart treatment when *r*_PBLC_ ≥ 5%, and then stop imatinib therapy at PBLC percentage level *r*_stop_ = 0.01% again. (A) Dynamics of PBLC ratio (*r*_PBLC_) for 20 patients. (B) Molecular RFS curves of patients. (C) LSPC percentage and TME index at the time of stopping imatinib treatment for the first time. Blue dots represent the data for TFR patients after imatinib discontinuation for the first time; cyan dots for patients with CML relapsed after the first imatinib discontinuation, but TFR after second imatinib discontinuation; red dots for relapsed patients after both the first and second imatinib discontinuations. (D) Cyan dots represent the data for TFR patients after imatinib discontinuation for the second time; red dots for relapsed patients after the second imatinib discontinuations.

### Importance of TME dynamics

In the proposed model, we assumed a positive feedback between leukemia stem cells and the tumor microenvironment, this positive feedback is crucial for the dichotomous response of patients between early relapse and long-term TFR after imatinib discontinuation. Nevertheless, we asked whether the interaction between leukemia stem cells and microenvironment is necessary to explain the clinical observations. To examine the effect of this interaction, we broke down the positive feedback, and assume a constant TME index in the model, and explored the patient responses after treatment discontinuation. Fig. 9 shows the time courses of PBLC ratios for patients with constant TME indices and stop treatment at MR4.0 (PBLC ratio *r*_PBLC_ < 0.01%).

**Figure 9:**
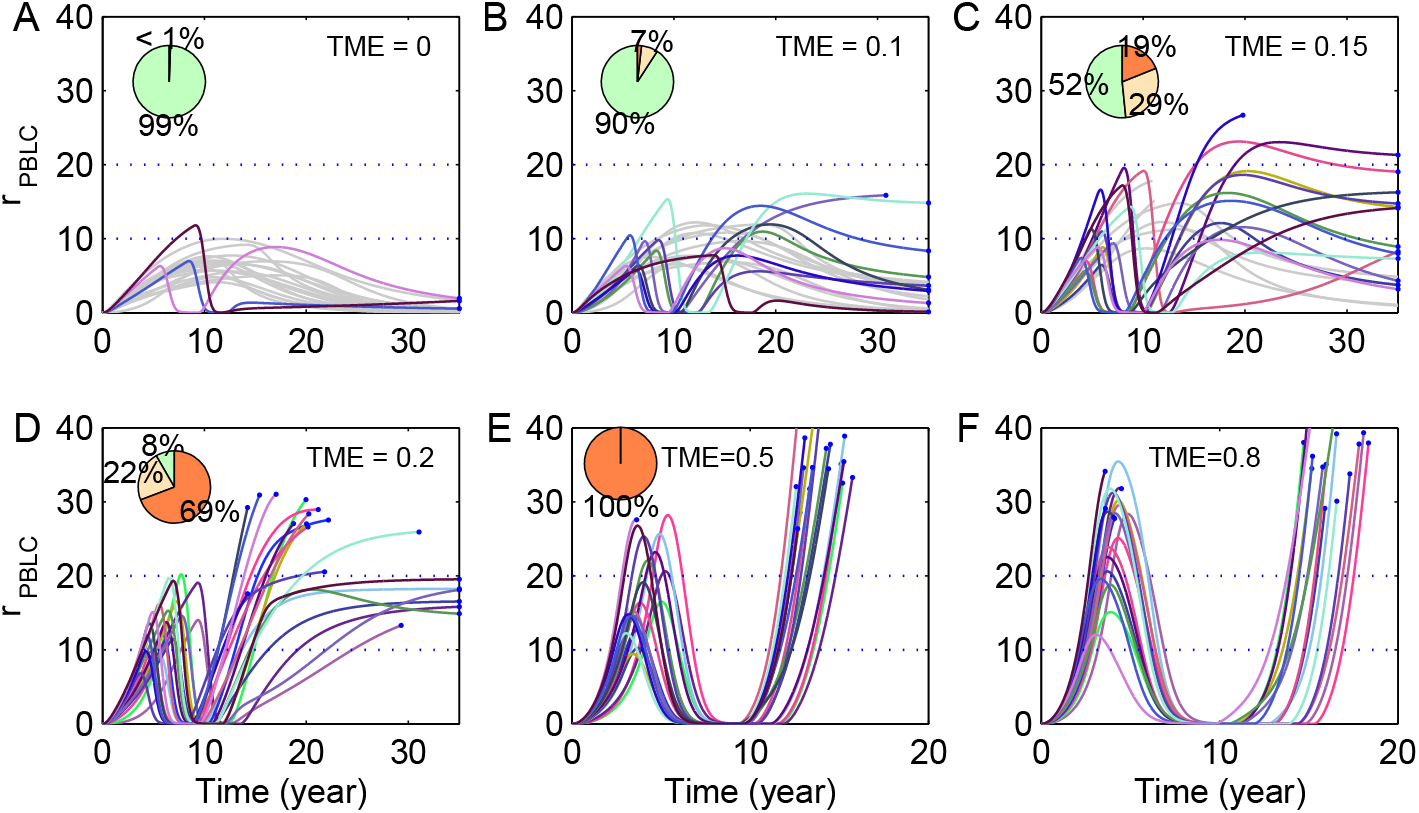
Evolution of PBLC percentage after treatment discontinuation based on the model with constant TME. (A)-(F): Evolution of PBLC percentage (*r*_PBLC_) for 20 patients with random initiation of TKI treatment when 5% < *r*_PBLC_ = 0.01% < 20%, and treatment discontinues when PBLC percentage level reach *r*_PBLC_ = 0.01%, with constant TME = 0, 0.1, 0.15, 0.2, 0.5 and 0.8, respectively. Pie plots in (A)-(E) show the percentages of patients under CP phase (green), AP phase (yellow), and CB phase (red) at 35 year, with simulation of 500 virtual patients, respectively. In (A)-(C), some patients automatically develop to CP phase without treatment, which are shown by gray lines.

As TME = 0, most patients remain at the CP phase and hence no treatment is required, and the others admitted with treatment show no molecular recurrence after treatment stop (Fig. 9A). When TME increases with TME = 0.1, 0.15, 0.2, the percentage of patients who developed AP and CB phases after treatment discontinuation obviously increase with the TME index (Fig. 9B-D). When TME becomes higher, with TME = 0.5, 0.8, all patients may progress to BC phase after treatment stop (Fig. 9B-D). Hence, we need to have TME < 0.5 while we assumed a constant TME in order to reach TFR after treatment discontinuation. Moreover, we show the distribution of *r*_PBLC_ from 1 to 7 years after treatment stop (Fig. 10), results show that dynamic change of TME can give rise to bimodal distribution of *r*_PBLC_ five years after treatment stop, however constant TME can only give unimodal distribution in *r*_PBLC_. These results suggest that changing TME due to the positive feedback between leukemia stem cells and microenvironment is essential for the dichotomous responses after treatment stop.

**Figure 10:**
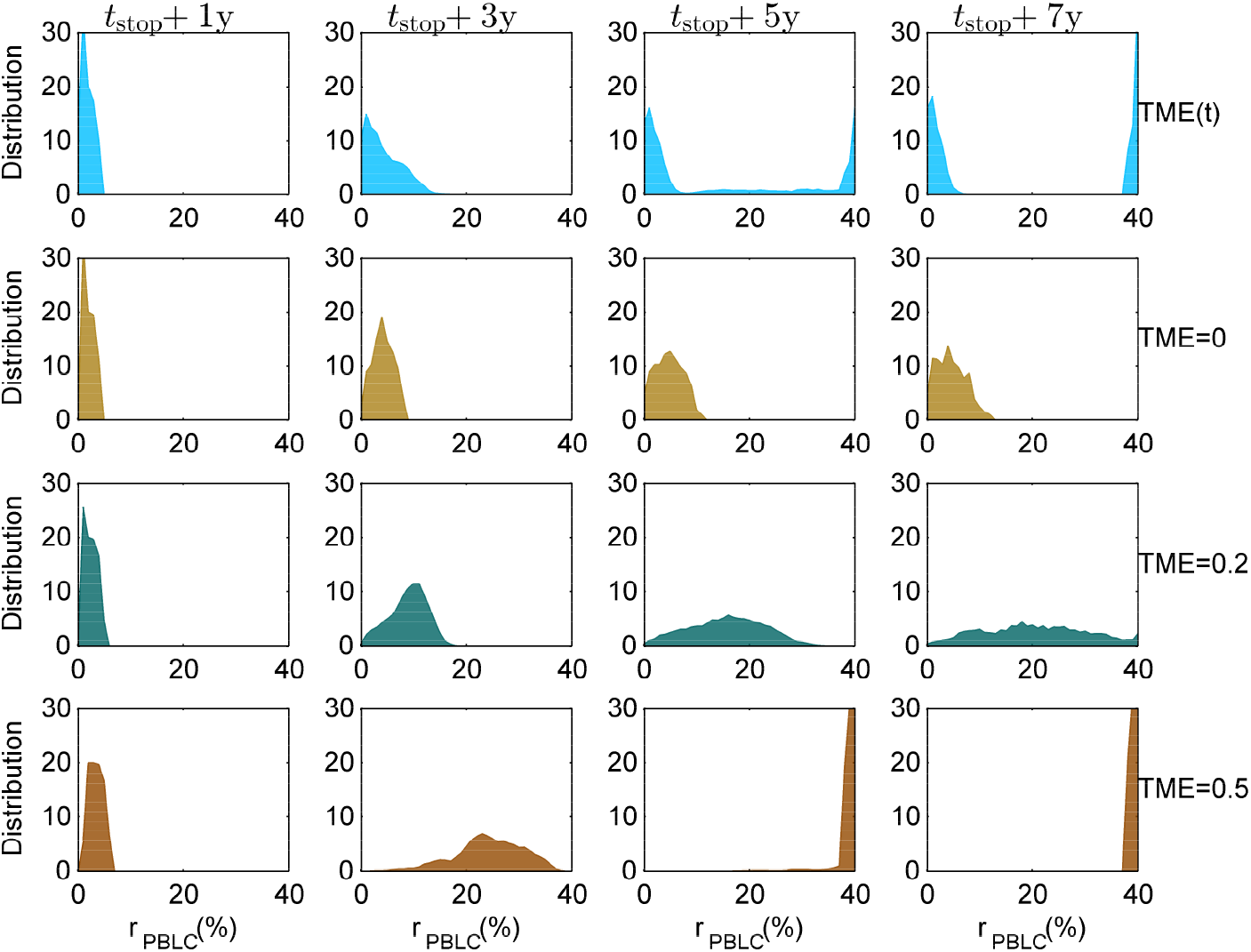
Evolution of PBLC ratio distribution after TKI treatment discontinuation for virtual patients. Four row panels show PBLC ratio distribution at 1, 3, 5, 7 years after TKI treatment discontinuation for virtual patients with dynamically changed TME based on the proposed stochastic differential equation model (TME(*t*)), or constant TME with TME = 0, 0.2, 0.5, respectively. Here, *t*_stop_ indicates the time point of TKI discontinuation. The mortality of the virtual patients are ignored in model simulations.

To further examine the the dynamics of CML evolution based on assumptions of changing TME and constant TME, we compare the time to reach AP phase and BC phase of patients after the emerge of BCR-ABL1 cells. The time spends to reaches AP phase for the patients with dynamically changed TME take values between those for patients with TME = 0.1 and TME = 0.15 (Fig. S7), whereas the time to BC phase for the patients with dynamically changed TME is close to that for patients with TME = 0.2 (Fig. S7). Hence, we would have TME < 0.2 in order to reasonably reproduce the progression of CML evolution. Furthermore, we compare the survival curve of virtual patients under continuous treatment, which show that only when the constant TME was taken as high as 0.8 can the model reproduce the overall survival rate of 80% under continuous treatment (Fig. S8). Nevertheless, we have seen that when TME = 0.8, all patients would undergo molecular recurrence after the TKI withdraw (Fig. 9F), which is inconsistent with experimental observations. These results suggest that the constant TME index is unlikely to reproduce CML evolution dynamics for both situations of no treatment and continuous treatment, and the hence dynamically changed TME index is important for CML evolution.

## Discussion

Chronic myeloid leukemia (CML) is one of a limited number of cancers that is critically dependent on a single genetic aberration. Tyrosine-kinase inhibitors (TKIs) have been a valuable treatment for CML patients. Effective treatment with TKIs leads to the majority of patients with CML to achieve a deep molecular response after ≥ 5 years of treatment. In recent years, international treatment guidelines for CML have incorporated recommendation of attempting TKI discontinuation toward the aim of treatment-free remission (TFR). Multiple clinical studies demonstrated that approximately 50% of patients with a sustained deep molecular response can discontinue the TKI and remain in TFR, and the majority of patients who have molecular relapse following discontinuation of TKIs do so within the first 6 months after treatment stop. Most relapsed patient can regain major molecular response after reinitiation of the original TKI, and some patients can reach second TFR after the first failed attempt and restart of TKI therapy. The biological and clinical factors that influence the outcome after TKI discontinuation are not well documented, predictors for TFR outcome are under investigation.

TKI treatment would not guarantee the elimination of all leukemia cells [40]. A study to identify immunophenotypically defined CML stem cells (CD34^+^CD38^*−*^CD26^+^) in patients with TFR showed that many patients had persistent cells of this type in the peripheral blood at 1 − 152 months after discontinuation of a TKI [21]. Thus, given a small number of residual leukemia cells upon deep molecular response, there is an apparent dichotomy in TKI-treated patients between early relapse and long-term TFR. This raises a sensible consequence that some factors other than minimal residual disease (MRD) may play essential roles for the TFR outcome.

In this study, we consider external cellular factors as the tumor microenvironment (TME) that is involved in the regulation of production and survival of leukemia stem and progenitor cells, and establish a stochastic mathematical model to investigate the dynamics of CML progression. Based on the model, both disease progressions without treatment and under imatinib therapy are quantitatively studied, and validated with criteria of the three CML evolution phases and clinical overall survival curves. The model predicts the dichotomous response of patients between early relapse and long-term TFR after imatinib discontinuation that is in agreement with clinical observations. Moreover, the dynamically changing TME index due to the feedback between leukemia stem cells and microenvironment is essential for the dichotomous responses. Based on model simulations, the remission process after imatinib treatment is not the identically reverse of CML evolution prior treatment in the phases of TME index and the ratio of leukemic cells in peripheral blood (PBLC). Moreover, the combination of change rates of TME index and bone marrow leukemia cells ratio 6 months prior imatinib therapy stopping provides a potential predictor of TFR outcome.

To our knowledge, the study reported here is the first try to explore the issue of TFR in CML patients through the mechanism-driven mathematical model. The interaction between tumor cells and microenvironment is highlighted in the proposed mechanism, and the state of TFR can be maintained by normal microenvironment even minimal residual leukemic cells may exist. The proposed model is essentially conceptual, and further extensions should be explored in order to include more details. For example, the heterogeneity of leukemic stem cells that may result in heterogeneous response to TKI treatments, the potential mutations to produce drug resistance, external perturbation to the microenvironment, and further characteristic description of the microenvironment. Clinically, how TME in the proposed model is associated with physiological index is yet to be investigated. Our study highlights the importance of microenvironment in TFR, effective methods to measure and control the microenvironment can play important roles for improving cancer therapy and the final goal of TFR.

## Materials and methods

### CML evolution

According to the model assumption in Fig. 1, dynamics of the cell population numbers of hematopoietic stem progenitor cells ([HSPC]), leukemia stem progenitor cells ([LSPC]), hematopoietic cells in peripheral blood ([PBHC]), leukemia cells in peripheral blood ([PBLC]), and the TME index (*Q*) are modeled with the following random differential equations (RDEs):

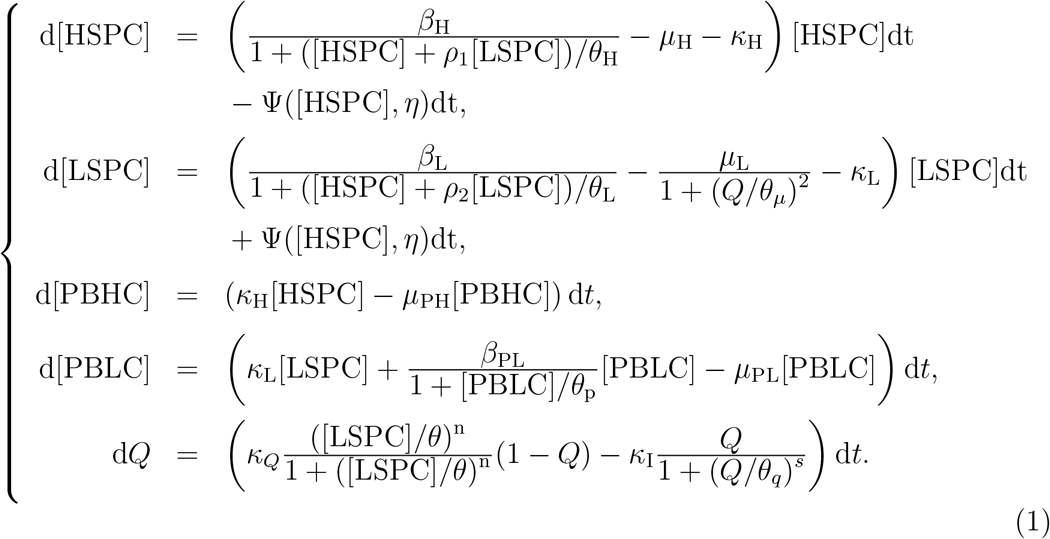

Here, *β*_H_, *β*_L_, and *β*_PL_ represent the maximum proliferation rates of HSPC, LSPC, and PBLC, respectively. Stem cell proliferation is subjected to the regulation of cytokines so that the proliferation rates are decreasing functions of cell numbers, which are formulated as Hill type functions 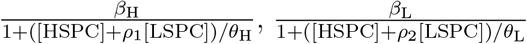, and 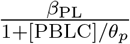, respectively. The coefficients *θ*_H_ and *θ*_L_ represent the difference between HSPC and LSPC in their proliferation ability that are related to the inhibition from HSPC and LSPC; *ρ*_1_ and *ρ*_2_ represent the relative strength of growth inhibitors released by LSPC with respect to HSPC. The parameters *µ*_*X*_ (*X* = H, L, PH) represent the apoptosis rates of corresponding cells, and *κ*_*X*_ (*X* = H, L) represent the differentiation rates of HSPC and LSPC from bone marrow to peripheral blood. The TME index *Q* is assumed to take values over [0, 1], with larger value of *Q* means more permissible for tumor cells survival, and the transition from normal to tumor microenvironment is positively regulated by tumor cells in bone marrow. We assumed that the microenvironment condition is essential for cell survival, and the apoptosis rate for LSPC 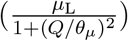 is a decreasing function of the TME index *Q*.

To model the origin of CML from health person, we assumed that the random generation of leukemia cells from normal hematopoietic progenitor stem cells is formulated as a Poisson random number Ψ([HSPC], *η*) that is dependent on the normal cell number [HSPC] and the rate *η* of BCR-ABL1 fusion gene formation. This gives the expectation number of LSPC generated from HSPC per unit time (day) as

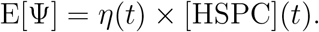

The probability of fusion gene formation is varied with time so that the rate *η* is a time dependent function. Moreover, the TME may also affect the generation and survival of cells with fusion genes. Hence, we wrote

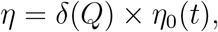

and

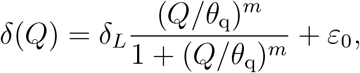

to represent the promotion of leukemia cell formation by TME. To model the progression of CML, we assumed that the time dependent rate *η*_0_ increases from 0 to a high level during disease progression, and decrease to a lower level latter on. Thus, phenomenologically, we write

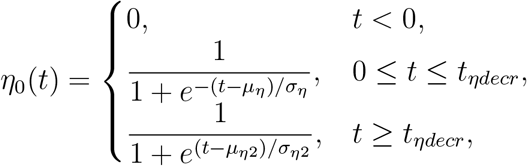

where *t*_*ηdecr*_ denotes the time when *η* starts to decrease.

### Imatinib treatment

To model imatinib treatment, we omitted the potential drug resistance, and assumed that imatinib can effectively result in the apoptosis of leukemia cells in both bone marrow and peripheral blood. Hence, there is an extra apoptosis rates to the leukemia cells so that *µ*_L_ and *µ*_PL_ in (1) are replaced by 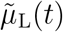 and 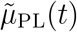 given below

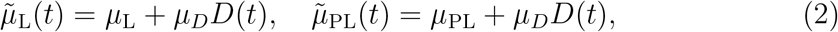

where *µ*_*D*_ represents the maximum cell death rate under imatinib treatment, and *D*(*t*) represent the drug delivery function. In model simulations, when we consider the situation of drug delivery from *t* = *t*_cr_, we set

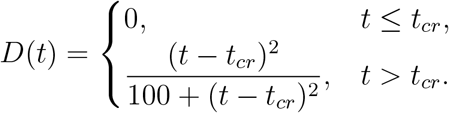

Here, the unit of the time *t* is day.

### Overall survival

To compare the overall survival curve with clinical data, we introduced a patient death rate that is dependent on the PBLC ratio. In simulations, we assumed

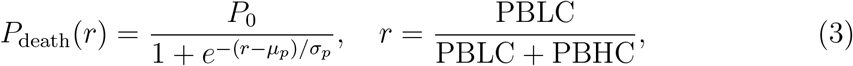

so that *P*_death_(*r*)Δ*t* gives the mortality probability of a patient with PBLC ratio *r* in a time window [*t, t* + Δ*t*].

### Parameter estimation

To estimate model parameters, we proposed restrictions in parameters that are biologically reasonable. First, leukemia cells are expected to have higher proliferation rate and lower death rate, and hence *β*_L_ > *β*_H_ and *µ*_L_ < *µ*_H_ [30]. We assumed that *β*_PL_ ≪ *β*_L_ since there is no strong evidence to support or against the division of leukemic cells in peripheral blood. Here, we took 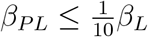. Moreover, the death rates in peripheral blood were assumed to be larger then those in the bone marrow, hence *µ*_PH_ > *µ*_H_, *µ*_PL_ > *µ*_L_. These assumptions give rise to the following restrictions in model parameters

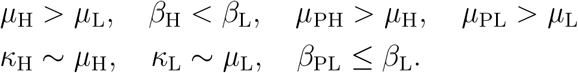

Here ∼ means the same order of magnitude.

We referred previous mathematical models of hematopoietic stem cells and CML progression dynamics for the overall magnitude of key parameters for the stem cell proliferation, differentiation, and apoptosis rates [30, 41], namely

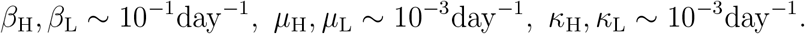

We performed random sampling of parameter values over a wide range of a parameter space, and identify the parameter values that produce the three phases of CML evolution dynamics (Fig. 2) and the overall survival curve without treatment (Fig. 3A). Key parameters are: *β*_H_ = 0.112day^*−*1^, *µ*_H_ = 0.0065day^*−*1^, *κ*_H_ = 0.0058day^*−*1^, *β*_L_ = 0.158day^*−*1^, *µ*_L_ = 0.0045day^*−*1^, *κ*_L_ = 0.0022day^*−*1^, *µ*_PH_ = 0.0088day^*−*1^, *β*_PL_ = 0.005day^*−*1^, *µ*_PL_ = 0.0060day^*−*1^. Next, we adjusted the parameters related to imatinib treatment to obtain the CML progression with imatinib therapy (Fig. A full list of parameter values are listed in Table S1.

### Virtual patients

To generate virtual patients in model simulations, we assumed heterogenous origins to induce BCR-ABL1 generation among different patients so that the rate function *η*_0_(*t*) is different among patients, which is defined by uniform random numbers (*µ*_*η*_, *σ*_*η*_, *µ*_*η*2_, *σ*_*η*2_). Moreover, we randomly selected the parameter values for the kinetic rates of proliferation (*β*_L_), apoptosis (*µ*_L_), and differentiation (*κ*_L_) of leukemia cells for different specific patients. Ranges of these parameters are listed in Table S2.

### Data Analysis

To quantify the progression of CML evolution, we analyzed the data set GSE4170 from GEO database [37, 36] that collected gene expression data of 91 CML patients in chronic phase (42 cases), accelerated phase (17 cases), and blast crisis (32 cases). We compared CD34 expression level in three phases (Fig. S1A), the average CD34 expression level increases with CML progression. Hence, we choose CD34 expression level as a marker to represent the disease age of CML starting from the emergence of BCR-ABL1 cells.

To define the disease age for individual patients, we assumed a linear dependence between the disease age and the CD34 expression level, and the coefficients can be different for patients at different phases. Hence, the disease age *T*_disease age_ (year) is formulated as

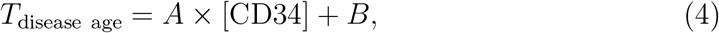

where (*A, B*) take values (5.0, 5.0) for chronic phase patients, (4.5, 6.5) for accelerated phase patients, and (4.0, 10) for blast crisis patients. Here [CD34] represents the normalized CD34 expression level of a patient. Dependence of the disease age with normalized CD34 level is shown at Fig. S1B.

**Figure S1:**
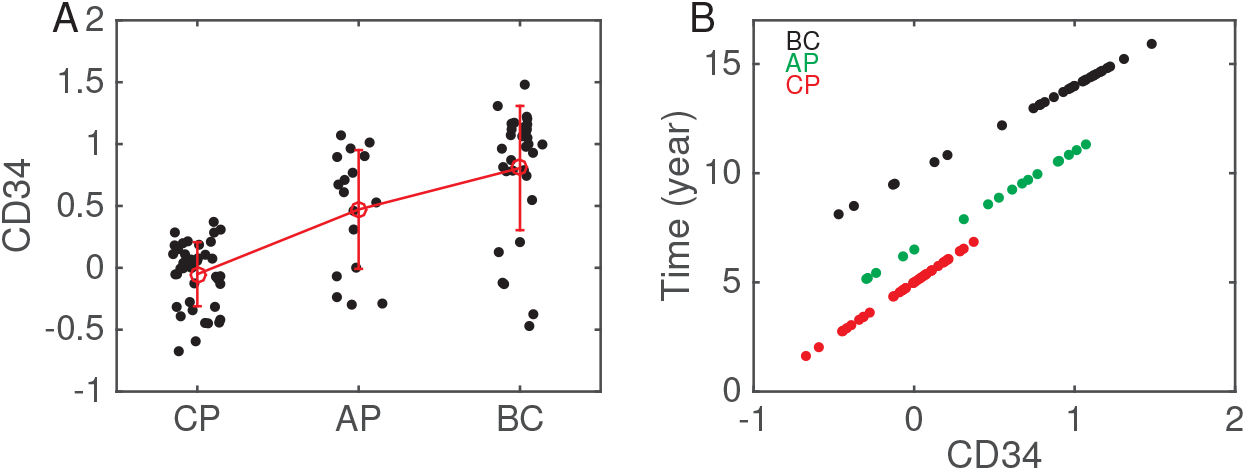
Data analysis of CML evolution. (A) CD34 expression level of patients in three phases of CML evolution: Chronic phase (CP), accelerated phase (AP), and blast crisis (BC). (B) Disease age defined by CD34 expression level. Red, green and black dots denote CP, AP and BC patients, respectively.

### Thresholds of TME index and LSPC for CML relapse

To obtain the threshold of TME index and LSPC number that can determined the patient responses after treatment discontinuation, we assumed that the tumor microenvironment in a quasi-steady state, then the TME index (*Q*) and LSPC ([LSPC]) satisfies

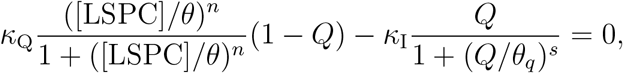

which gives

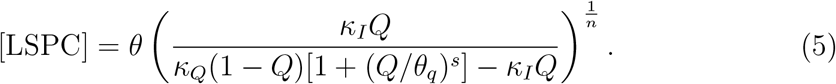

The critical zone for LSPC and TME index in Fig. 4 is given by this formula, where *κ*_*Q*_ varies over the range 0.006 ≤ *κ*_*Q*_ ≤ 0.0078 and the other parameters are chosen as in Table S1.

## Acknowledgement

This work is supported by the National Natural Science Foundation of China (No. 11831015 and No. 12171478).

## Author contributions

Jinzhi Lei and H. Z. designed research; Xiulan Lai, X. J. performed research and analyzed data; Xiulan Lai, X. J., H. Z., and J. L. wrote the paper.

## Supplemental Figures

**Figure S2:**
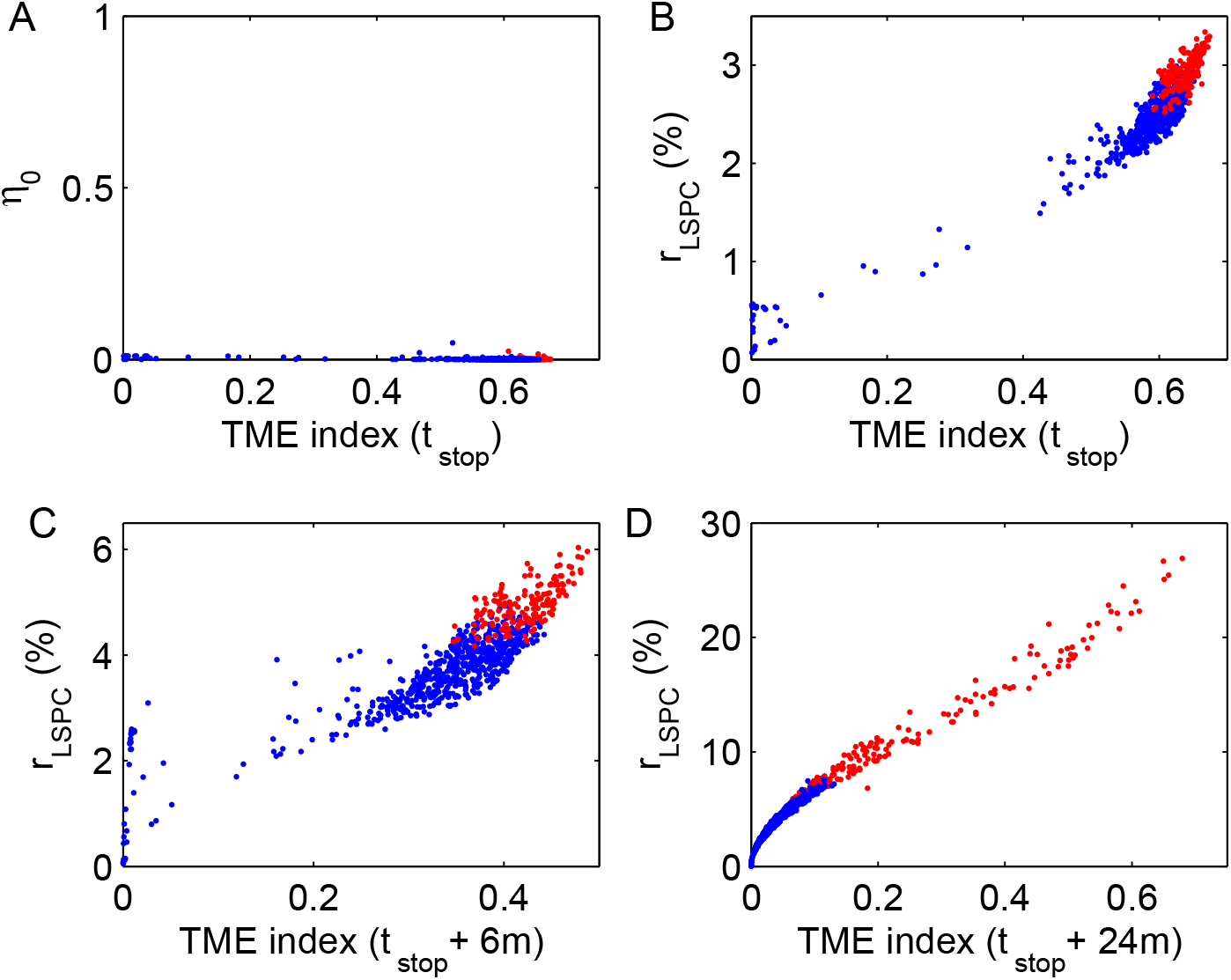
Classification of patients by TME index and LSPC percentage after TKI discontinuation under a certain pattern of mutation *η*_0_. (A) TME index and mutation rate *η*_0_ at the time of TKI discontinuation. (B) LSPC percentage and TME index at the time of TKI discontinuation. (C)-(D) LSPC percentage and TME index at the time of 6 months and 12 months after stopping treatment, respectively. Here the treatment is stopped when PBLC percentage level reach *r*_PBLC_ = 0.01%. Red dots for relapsed patients, and blue dots for TFR patients. Here, we fixed *µ*_*η*_2 = 9 year so that *η*_0_ becomes nearly zero at the time of TKI stop.

**Figure S3:**
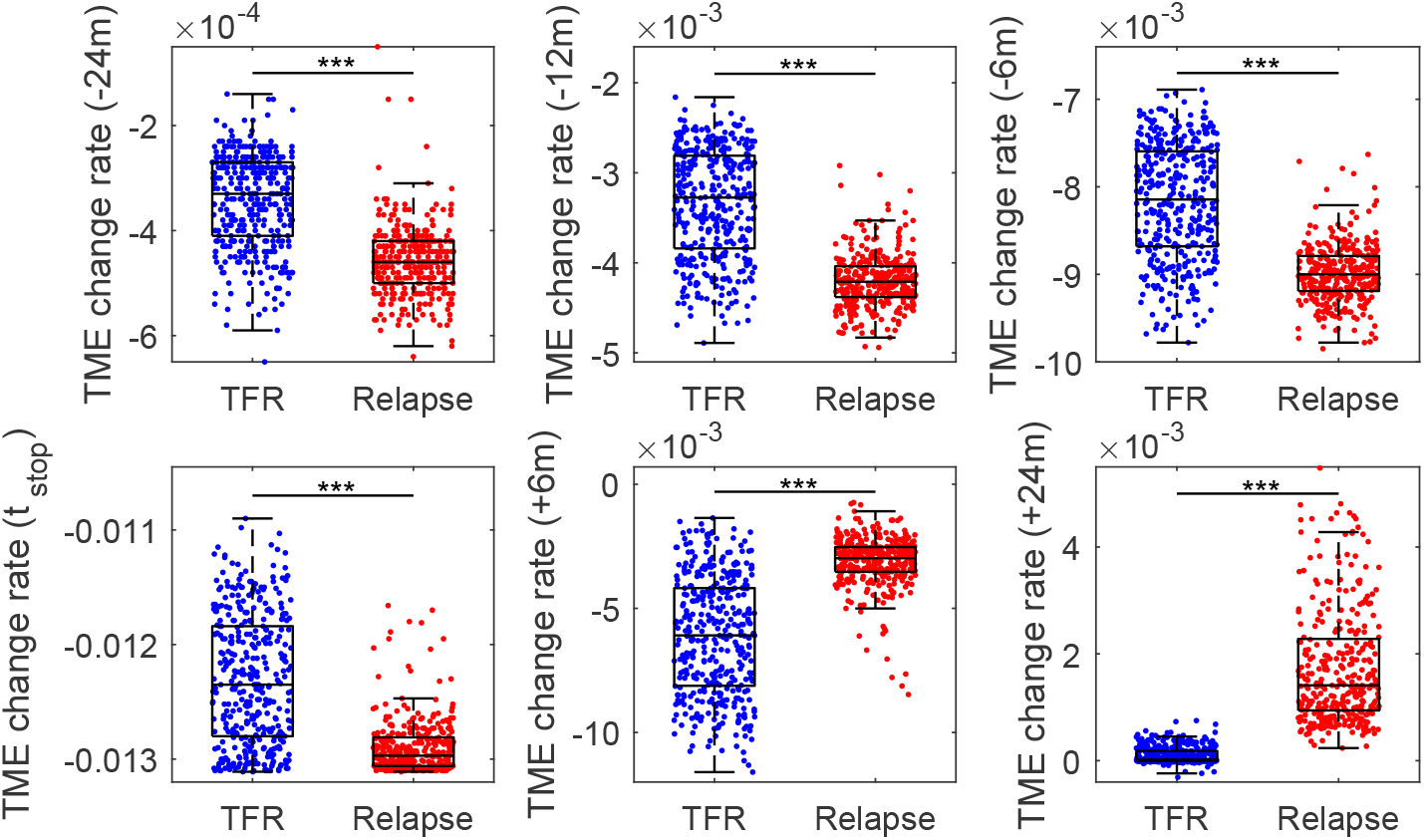
Comparison of TME change rates for TFR patients and relapsed patients. TME change rates for TFR patients and relapsed patients at 2 years (−24m), 1 year (−12m), 6 months (− 6m) before TKI stop, at the time TKI stop (*t*_stop_), at 6 months (+6m) and 2 years (+24m) after TKI stop. P-value: *** *p* < 0.001.

**Figure S4:**
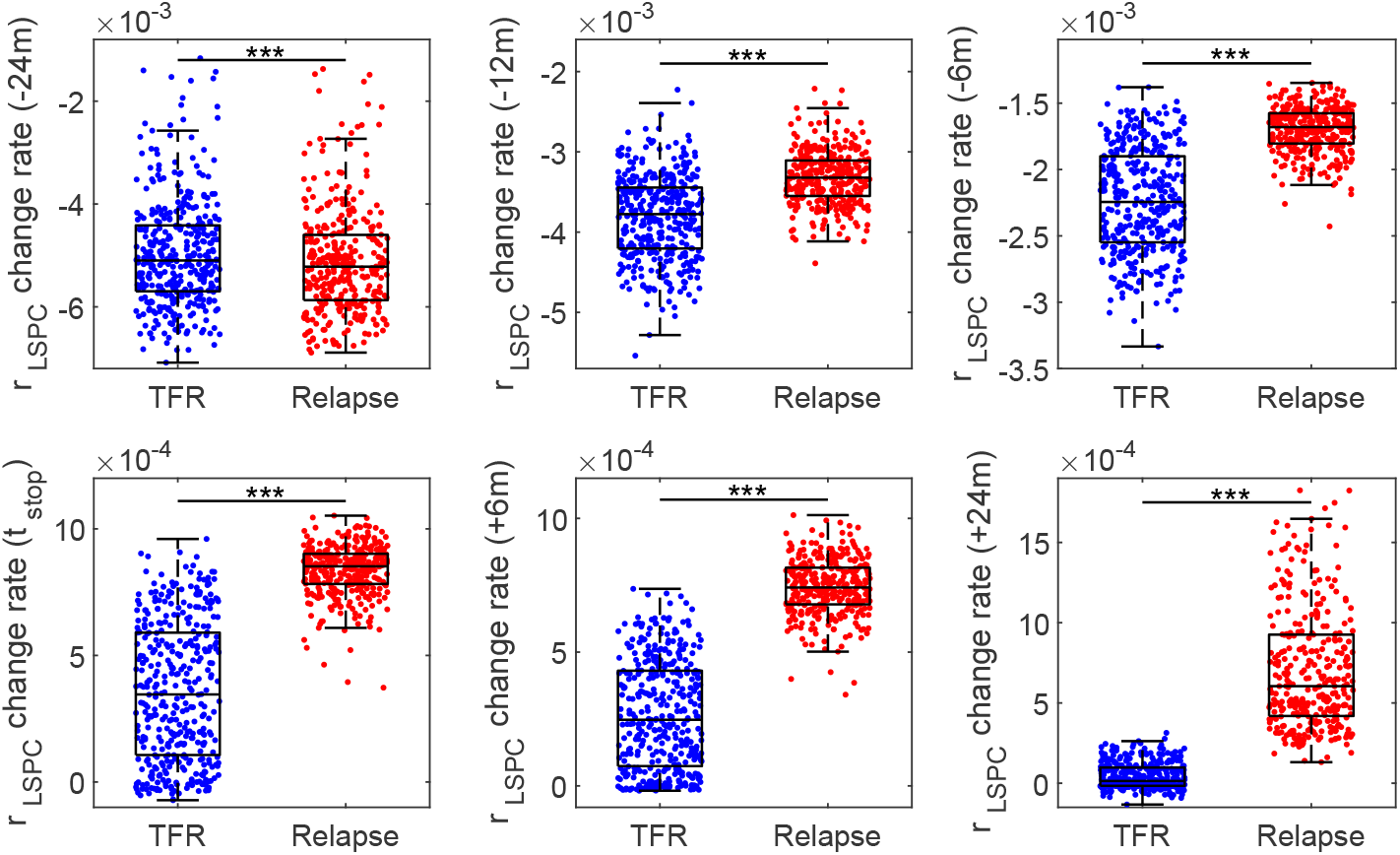
Comparison of change rates of bone marrow leukemia cells ratio (*r*_LSPC_) for TFR patients and relapsed patients. Change rates of bone marrow leukemia cells ratio for TFR patients and relapsed patients at 2 years (−24m), 1 year (−12m), 6 months (−6m) before TKI stop, at the time TKI stop (*t*_stop_), and at 6 months (+6m) and 2 years (+24m) after TKI stop. P-value: *** *p* < 0.001

**Figure S5:**
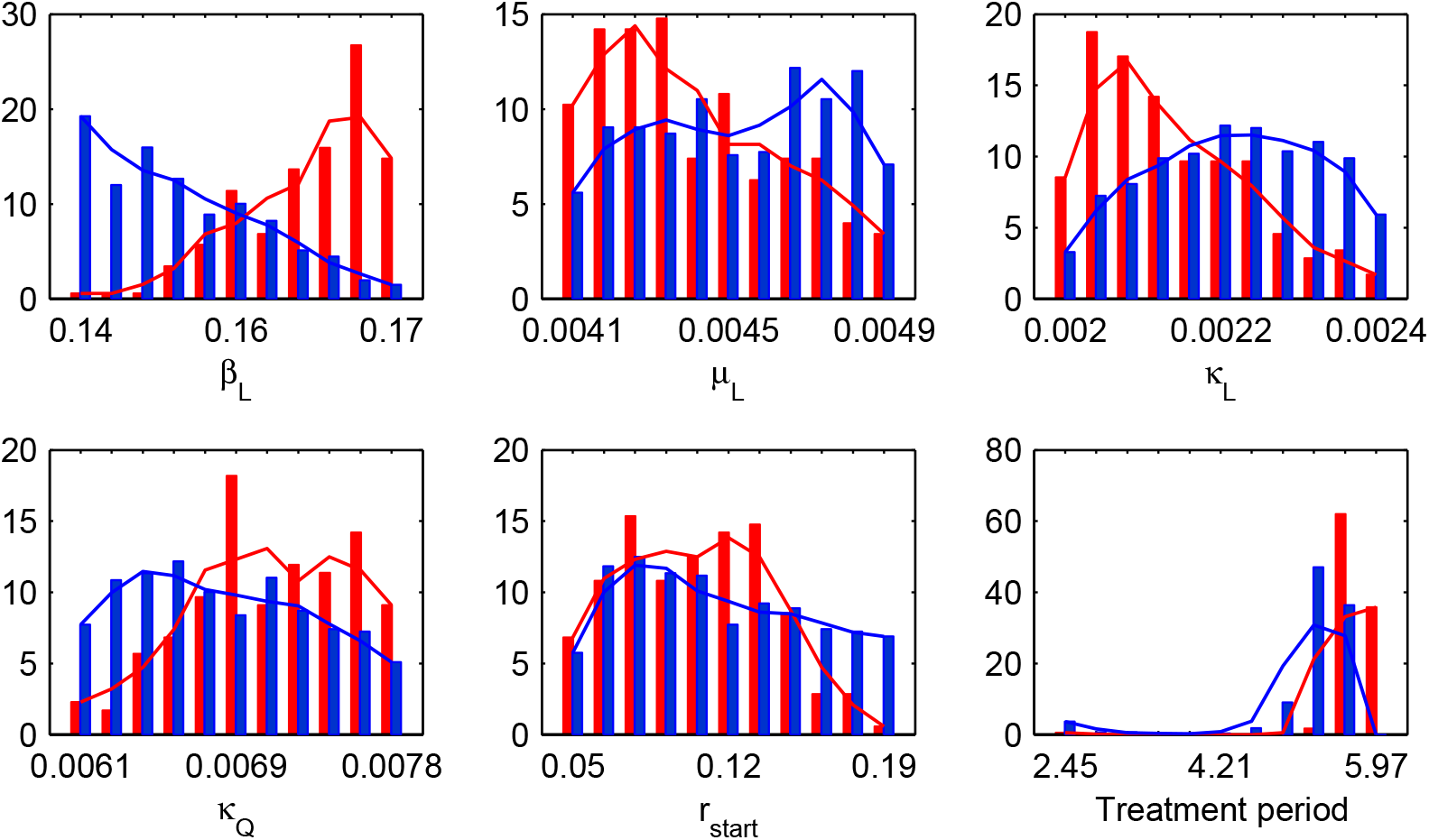
Distribution of parameter values for relapsed and TFR patients. The parameter value distributions of *β*_*L*_, *µ*_*L*_, *κ*_*L*_ and *κ*_*Q*_, and the distributions of TKI initiation phase (PBLC percentage at TKI initiation *r*_start_) and the treatment period for relapsed patients (red) and TFR patients (blue). Here, mutation rate *η*_0_ was set to the same as in Fig. S2.

**Figure S6:**
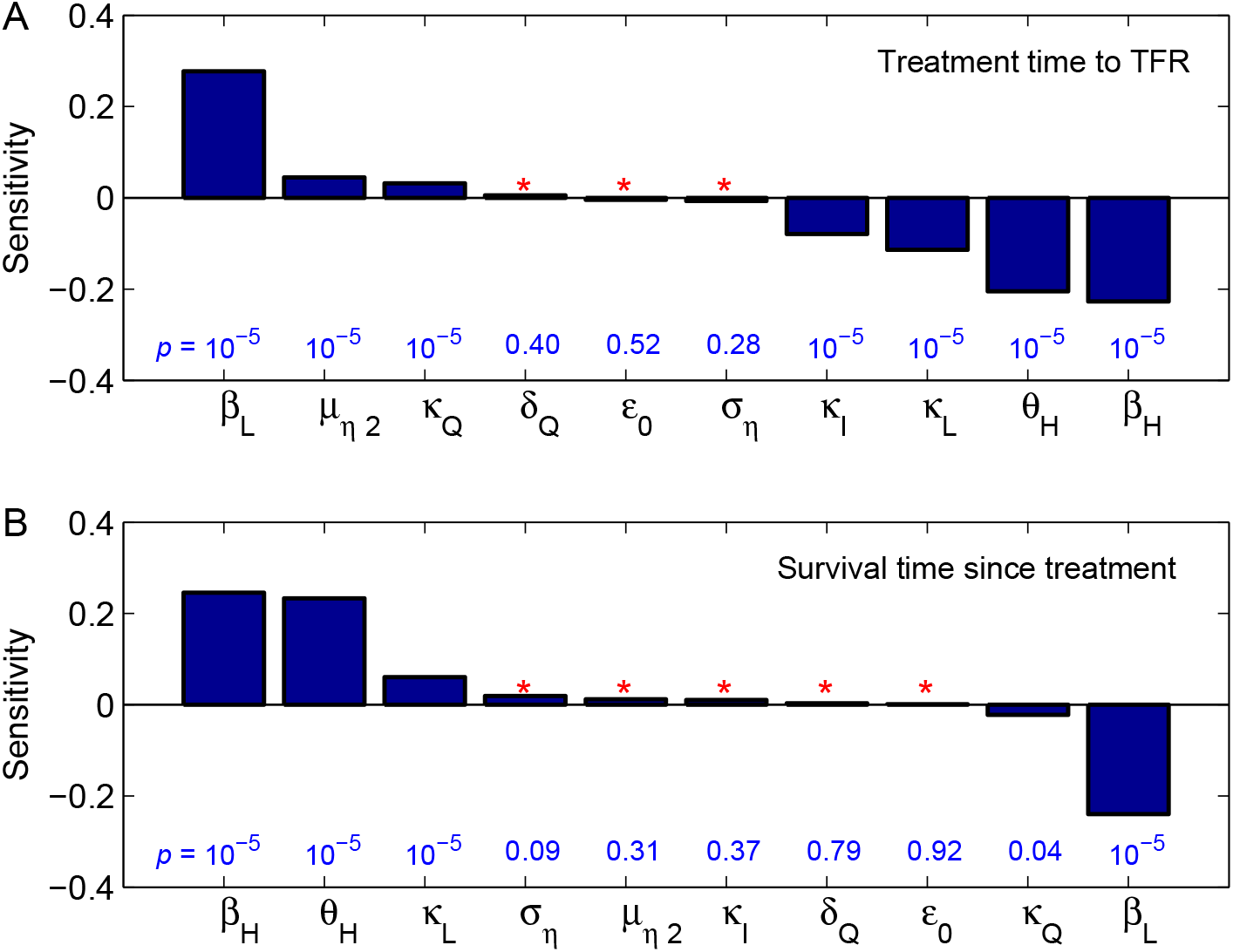
Sensitivity analysis. (A) Pearson’s correlation coefficients of the treatment time to reach TFR (for TFR patients) with the parameters *β*_*H*_, *θ*_*H*_, *β*_*L*_, *κ*_*L*_, *κ*_*Q*_, *κ*_*I*_, *σ*_*η*_, *µ*_*η*2_, *δ*_*Q*_, and *ε*_0_. (B) Pearson’s correlation coefficients of survival time after treatment (for dead patients) with the parameters *β*_*H*_, *θ*_*H*_, *β*_*L*_, *κ*_*L*_, *κ*_*Q*_, *κ*_*I*_, *σ*_*η*_, *µ*_*η*2_, *δ*_*Q*_, and *ε*_0_. The data values in blue indicate the corresponding p-values, where 10^*−*5^ represents p-values equal to or less than 10^*−*5^. The red star signs indicates statistically insignificant (with p-values greater than 0.05).

**Figure S7:**
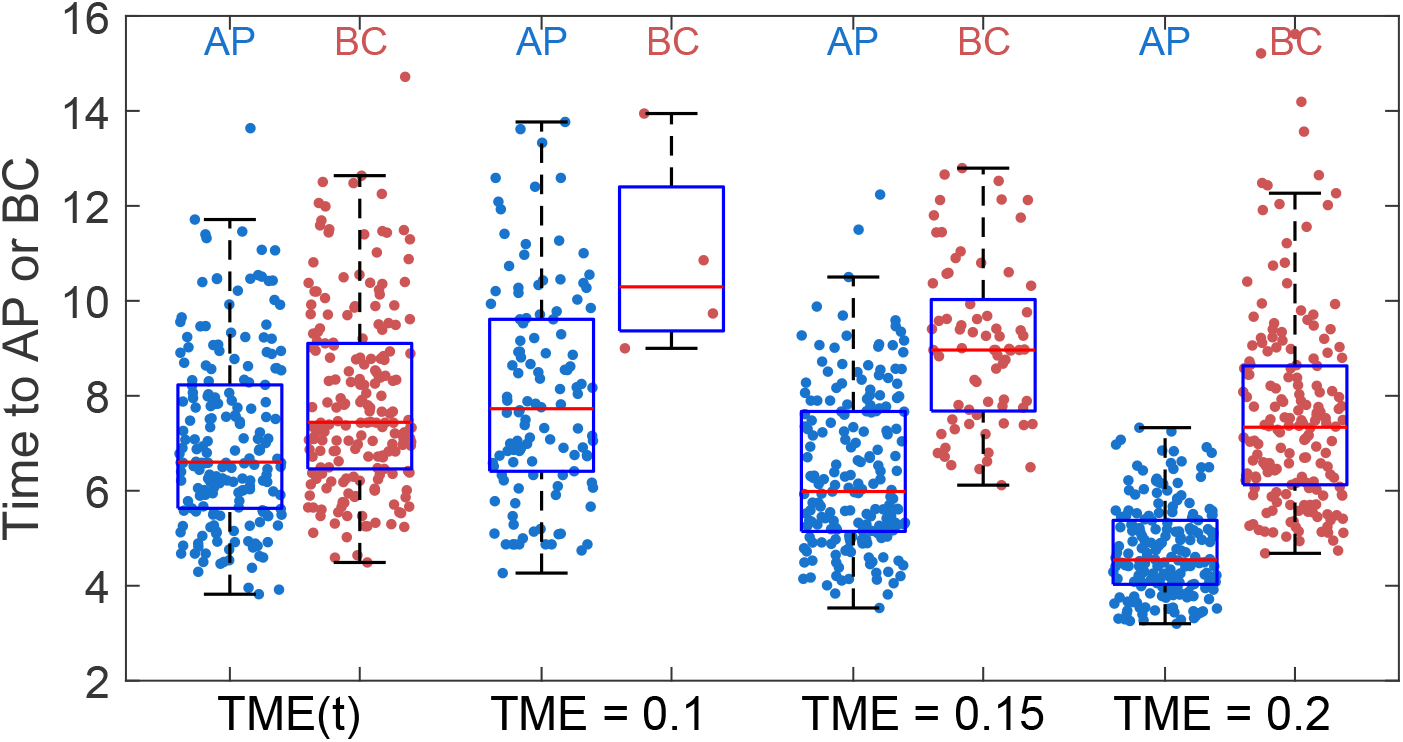
**Comparison of the time to AP and BC phases for models with dynamically changed TME (**MET(*t*)) **or constant TME indices with** TME = 0.1, 0.15 **and** 0.2, **respectively**.

**Figure S8:**
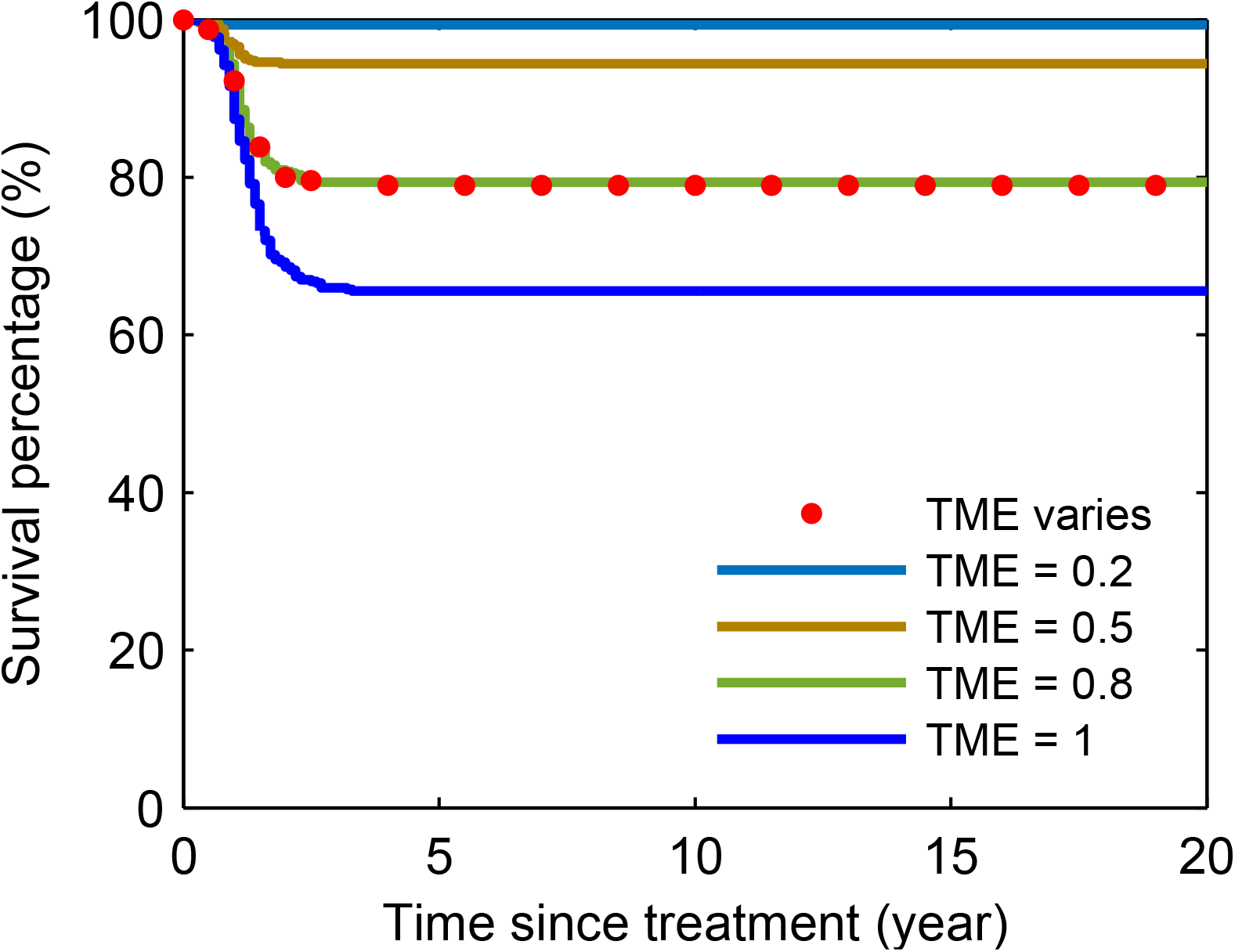
Survival curves of virtual patients after continuous TKI treatment. Dotted line shows the survival curve of virtual patients with dynamically changed TME based on the proposed stochastic differential equation model. Solid lines are survival curves obtained from constant TME = 0, 0.1, 0.15 and 0.2, respectively.

**Table S1:** Parameter values of the model.

| Parameter | Description | Value | Unit |
| --- | --- | --- | --- |
| $\beta_H$ | Basic proliferation rate of HSC | 0.112 | day<sup>-1</sup> |
| $\theta_H$ | Effective concentration for HSPC inhibition by HSPC and LSPC | $1.8 \times 10^6$ | cells/kg |
| $\rho_1$ | Ratio factor | 0.75 | - |
| $\mu_H$ | Death rate of HSPC | 0.0065 | day<sup>-1</sup> |
| $\kappa_H$ | Differentiation rate of HSPC | 0.0058 | day<sup>-1</sup> |
| $\mu_{PH}$ | Death rate of PBLC | 0.0088 | day<sup>-1</sup> |
| $\beta_L$ | Basic proliferation rate of LSPC | 0.158 | day<sup>-1</sup> |
| $\kappa_L$ | Differentiation rate of LSPC | 0.0022 | day<sup>-1</sup> |
| $\mu_L$ | Death rate of LSPC | 0.0045 | day<sup>-1</sup> |
| $\theta_L$ | Effective concentration for LSPC inhibition by HSPC and LSPC | $0.57 \times 10^6$ | cells/kg |
| $\rho_2$ | Ratio factor | 1.25 | - |
| $\theta_\mu$ | Effective concentration for inhibition of LSPC death by TME index $Q$ | 0.25 | cells/kg |
| $\beta_{PL}$ | Maximum proliferation rate of PBLC | 0.005 | day<sup>-1</sup> |
| $\theta_p$ | Effective concentration for PBLC proliferation regulation | $1.0 \times 10^5$ | cells/kg |
| $\mu_{PL}$ | Death rate of PBLC | 0.006 | day<sup>-1</sup> |
| $\kappa_Q$ | Rate of transformation from NME to TME | 0.0069 | day<sup>-1</sup> |
| $\kappa_I$ | Rate of restoring NME from TME | 0.015 | day<sup>-1</sup> |
| $\theta$ | Effective concentration of LSPC in promoting NME to TME transition | $2.0 \times 10^6$ | day<sup>-1</sup> |
| $\delta_L$ | Maximum production rate of LSPC from HSPC | 0.0043 | day<sup>-1</sup> |
| $\varepsilon_0$ | Fraction of residual transformation rate of HSPC to LSPC | 0.027 | day<sup>-1</sup> |
| $\theta_q$ | Effective TME in inhibition of TME depredation | 0.25 | cells/kg |
| $m$ | Hill coefficient | 2 | - |
| $n$ | Hill coefficient | 2 | - |
| $s$ | Hill coefficient | 2 | - |
| $\mu_D$ | Maximum TKI drug effect in promotion of LSPC apoptosis | 0.45 | day<sup>-1</sup> |
| $P_0$ | Maximum value of death probability of a patient | 1.0 | day<sup>-1</sup> |
| $\mu_P$ | Parameter related to the death probability of a patient | 0.5 | - |
| $\sigma_P$ | Parameter related to the death probability of a patient | 0.03 | - |

**Table S2:** Ranges of parameter values in generating the virtual patients.

| Parameters | Description | Range | Unit |
| --- | --- | --- | --- |
| $\beta_L$ | Basic proliferation rate of LSPC | [0.1422, 0.1738] | day<sup>-1</sup> |
| $\mu_L$ | Apoptosis rate of LSPC | [0.004, 0.005] | day<sup>-1</sup> |
| $\kappa_L$ | Differentiation rate of LSPC | [0.002, 0.0024] | day<sup>-1</sup> |
| $\kappa_Q$ | Rate of transformation from NME to TME | [0.06, 0.078] | day<sup>-1</sup> |
| $\mu_\eta$ | Parameter relate to BCR-ABL1 occurrence $\eta_0(t)$ | [0, 5] | year |
| $\sigma_\eta$ | Parameter relate to BCR-ABL1 occurrence $\eta_0(t)$ | [1, 4] | year |
| $\mu_{\eta 2}$ | Parameter related to BCR-ABL1 occurrence $\eta_0(t)$ | [9, 22] | year |
| $\sigma_{\eta 2}$ | Parameter related to BCR-ABL1 occurrence $\eta_0(t)$ | [0.2, 3] | year |

